# A Comparative Review of Deep Learning Methods for RNA Tertiary Structure Prediction

**DOI:** 10.1101/2024.11.27.625779

**Authors:** Ivona Martinović, Tin Vlašić, Yang Li, Bryan Hooi, Yang Zhang, Mile Šikić

## Abstract

Several deep learning-based tools for RNA 3D structure prediction have recently emerged, including DRfold, DeepFoldRNA, RhoFold, RoseTTAFoldNA, trRosettaRNA, and AlphaFold 3. In this study, we systematically evaluate these six models on three datasets: RNA Puzzles, CASP15 RNA targets, and a newly generated dataset of sequentially distinct RNAs, which serves as a benchmark for generalization capabilities. To ensure a robust evaluation, we also introduce a fourth, more stringent dataset that contains both sequentially and structurally distinct RNAs. We observed that each model predicts the best structure for certain RNAs, and evaluated whether commonly used scoring functions, Rosetta score and ARES, can reliably identify the most accurate structure from the predictions. Finally, since many RNA chains in the Protein Data Bank are part of complexes, we compare the performance of RoseTTAFoldNA and AlphaFold 3 in predicting RNA structures within complexes versus isolated RNA chains extracted from these complexes. This comprehensive evaluation highlights the strengths and limitations of current deep learning-based tools and provides valuable insights for advancing RNA 3D structure prediction.

## 1 Introduction

RNA molecules play key roles in various biological processes, ranging from gene regulation to catalysis [1]. Their functions are closely tied to their 3D structures, making accurate RNA structure prediction essential for understanding RNA biology and for applications in drug discovery and synthetic biology [2, 3, 4]. Experimental methods such as X-ray crystallography and cryo-electron microscopy are often costly and time-consuming, highlighting the need for computational approaches.

The field of protein structure prediction witnessed a paradigm shift with the introduction of AlphaFold2, which achieved remarkable success in predicting protein 3D structures with near-experimental accuracy. Inspired by AlphaFold2’s success, similar approaches have been adapted to RNA 3D structure prediction. The models we examine in this study, DRfold [5], DeepFoldRNA [6], RhoFold [7], RoseTTAFoldNA [8], trRosettaRNA [9], and AlphaFold 3 [10], build upon deep learning strategies originally developed for proteins and apply them to the unique challenges posed by RNA molecules.

While the adoption of deep learning has advanced the field, it is important to note that many machine learning-based tools for RNA structure prediction, such as those relying on probabilistic graphical models or feature-based methods, predate these advances. Given that each of these deep learning tools has been evaluated primarily against traditional methods, our work concentrates on comparing them directly against each other, with an additional focus on their ability to generalize to unseen, structurally novel RNA sequences.

The significant interest in RNA 3D structure prediction is evident from the growing number of recently published reviews [11, 12, 13, 14] and comparative studies [15, 16, 17, 18, 19]. However, while several studies have compared RNA 3D structure prediction tools, only a few include AlphaFold 3 alongside the other tools, and none evaluated the locally installed version of AlphaFold 3. Instead, they rely on the web server. The studies often fail to compare all state-of-the-art deep learning-based models on a consistent dataset. Moreover, these comparisons are often conducted on datasets that may overlap with the tools’ training data. As a result, they do not effectively evaluate the ability of these tools to generalize to unseen data. Similarly, many benchmarks rely on relatively small datasets, often with fewer than 40 RNAs, which limits the significance of the results. Furthermore, many studies fail to systematically assess sequence and especially structural similarity between the evaluation and training datasets, which is crucial for understanding true generalization performance.

In this study, we aim to address these limitations by comprehensively comparing six deep learning-based RNA 3D structure prediction tools across four datasets. Our key contributions are:

- **Comprehensive comparison**: We benchmark six tools, including locally installed AlphaFold 3, across three datasets: RNA Puzzles [20, 21, 22, 23], CASP15 RNA targets [24, 25], and a novel dataset of 84 RNAs published in the Protein Data Bank (PDB) [26] after 13 January 2023, referred to as Dataset 3. This dataset, larger than those used in prior studies, ensures no overlap with the models’ training data. Comprising RNAs that are sequentially distinct both among themselves and compared to all RNAs published prior to this cutoff date, Dataset 3 enables us to assess models’ generalization capabilities.
- **Evaluation of generalization**: Beyond Dataset 3, we created a fourth, more stringent dataset by selecting RNAs from Dataset 3 that are both sequentially and structurally distinct from RNAs in the training datasets, allowing us to rigorously test the models’ ability to generalize to highly dissimilar RNAs, presenting a more challenging and realistic test of their performance.
- **Evaluation of scoring function effectiveness**: We assess commonly used scoring functions, Rosetta score [27] and ARES [28], to determine their effectiveness in identifying the most accurate predicted structures. This approach is motivated by our observation that all models predicted the best structure for at least some RNAs. By incorporating scoring functions, we aim to explore what is the most we can get out of the benchmarked prediction tools.
- **Context-dependent performance comparison**: For RNAs from Dataset 3 that are a part of a complex in the PDB, we predict the entire complex using RoseTTAFoldNA and AlphaFold 3. We then compare these predictions to the structures obtained by modeling the isolated RNA chains.

This study is the first to combine the evaluation of structural similarity, scoring functions, and context-dependent predictions into a unified benchmark, offering a comprehensive assessment of state-of-the-art RNA 3D structure prediction tools. Our findings provide valuable insights into the strengths and limitations of current methods, as well as guidance for future improvements in RNA 3D structure prediction.

Since this study focuses on RNA tertiary or 3D structure prediction, any mention of structure prediction throughout refers specifically to RNA 3D structures unless otherwise specified.

## 2 Methods

This section briefly describes the benchmarked tools, datasets used, and metrics calculated for the evaluation.

### 2.1 Tools

This benchmark evaluates six deep learning-based RNA structure prediction models, each bringing certain unique approaches to the complex task of RNA tertiary structure modeling, which will be described in this subsection. The characteristics of these models are summarized in Table 1.

**Table 1.** Properties of deep learning-based RNA 3D structure prediction tools. *Abbreviations used: SS = secondary structure, MSA = multiple sequence alignment*.

| Tool | Input SS | Input MSA | End-to-end | Post-processing | Uniqueness |
| --- | --- | --- | --- | --- | --- |
| DRfold | RNAfold [29],<br>PETfold [30] | no | yes | optimization<br>using end-to-end<br>model +<br>geometric<br>restraints | predicts both<br>end-to-end<br>models and<br>geometric<br>restraints |
| DeepFoldRNA | PETfold (from<br>MSA) | rMSA [31] | no | refinement using<br>SimRNA [32] | × |
| RhoFold | no | rMSA, Infernal<br>[33] | yes | relaxation | LM-based<br>embeddings [34] |
| RoseTTAFoldNA | no | lite rMSA | yes | no | predicts RNA<br>and complexes |
| trRosettaRNA | SPOT-RNA [35] | rMSA, Infernal | no | no | × |
| AlphaFold 3 | no | nhmmer &<br>hmmalign | yes | no | diffusion-based<br>architecture,<br>works with<br>atoms, predicts<br>complexes |

**DRfold** takes sequence and secondary structure as inputs and generates embeddings using RNA transformer blocks. Uniquely, DRfold is the only tool that relies solely on the sequence without requiring a multiple sequence alignment (MSA). These embeddings are then fed into a structure module and a geometric module. The structure module predicts nucleotide-wise frames to build full 3D structures in an end-to-end fashion, while the geometric module estimates pairwise nucleotide geometries. By combining outputs from both modules to define energy terms, DRfold gets its final prediction through energy minimization using the L-BFGS algorithm.

**DeepFoldRNA** processes MSA and secondary structure representations through self-attention layers and linear projections to generate embeddings. These embeddings are then passed to linear layers that predict distance and orientation restraints between nucleotide pairs. Using these restraints to define an energy function, DeepFoldRNA optimizes the structure through L-BFGS minimization to produce the final RNA conformation.

**RhoFold** (also called E2Efold-3D) first processes MSA using an RNA language model (RNA-FM [34]). These representations go through E2Eformer/Rhoformer, which is based on an attention mechanism, similar to the previous two models, where the key is communication between MSA representation and pair representations. This module is shallower compared to other modules for sequence embedding updates (eg. DRfold has 48 RNA transformer blocks, DeepFoldRNA 48 MSA Transformer blocks, while Rhofold has only 4 Rhoformer blocks). After obtaining these embeddings, they go into 8 layers of structure module, which produce full atom structure.

**RoseTTAFoldNA** is an adaptation of RoseTTAFold [36, 37], initially designed for protein modeling, extended to predict structures of nucleic acids. This tool can model not only isolated RNA molecules but also more complex structures that include various combinations of proteins, DNA, and RNA. Distinct from other RNA-focused tools, RoseTTAFoldNA uses a three-track architecture: a sequence track that processes MSA inputs, a 2D geometric track capturing base-pair and stacking interactions, and a 3D track that incorporates either structural templates or iteratively refined nucleic acid coordinates. Together, these tracks enable a multi-level representation of the nucleic acid, progressively building up spatial detail and structural accuracy. In the final stage, RoseTTAFoldNA employs a structure refinement module based on the SE(3)-Transformer [38], which leverages symmetry-aware transformations to refine the predicted 3D coordinates, ensuring more accurate and biologically relevant conformations.

Similarly, **trRosettaRNA** builds on the idea from trRosetta [39], a tool initially developed for protein folding through distance and orientation predictions between residue pairs. This adaptation extends those principles to RNA by predicting distances and angles between nucleotide pairs, thus capturing key geometric relationships within RNA structures. Deep learning is applied to derive pairwise nucleotide geometries from MSAs, which then inform energy terms. These energy terms are combined with those from the PyRosetta suite, and energy minimization is conducted to refine the RNA model and achieve the final conformation.

**AlphaFold 3** builds directly on the AlphaFold family [40, 41, 42], which transformed protein structure prediction with the breakthrough of AlphaFold2. AlphaFold 3 extends this architecture and methodology to accommodate various molecules, including RNAs. One of the primary adjustments involves handling atoms directly rather than using frames to represent residues, allowing it to manage different residue types such as amino acids and nucleotides, as well as small molecules and ions. Additionally, AlphaFold 3 incorporates a diffusion-based structure module instead of the previous Structure module. It can predict RNA structures both as isolated chains and as part of larger RNA-protein complexes.

All models, including AlphaFold 3, were installed and executed locally on an NVIDIA H100 GPU card. AlphaFold 3 had a seed for each job set to 1. The only exception was for predicting complexes with AlphaFold 3, where the AlphaFold 3 web server was used instead due to the long execution times and resources required for such large complexes.

### 2.2 Datasets

In this benchmark, we used four datasets, all described in this section.

**Dataset 1** consists of 37 RNA Puzzles from the RNA-Puzzles initiative, a widely-used benchmark. To prevent overlap with Dataset 2, Puzzles 35 and 36, which were CASP15 targets, were excluded. Since most of these Puzzles were published in the Protein Data Bank (PDB) [26], some of them may be part of the training datasets for some of the tools. Even though this dataset may not fully show the generalization capabilities of each model, we included it for its historical value and to demonstrate performance on typical tasks. This approach reflects practical scenarios where, for instance, a structural biologist seeking to predict the structure of a common RNA, such as tRNA, may prioritize reliable results over concerns about potential overlap with training data.

**Dataset 2**. Historically, CASP primarily focused on assessing the accuracy of protein structure prediction methods. However, the 15th edition marked a significant milestone by including RNAs for the first time [24, 25]. This integration highlights the growing importance of RNA structure prediction in molecular biology and biotechnology. CASP15 presented 12 RNA targets, containing both natural and synthetic RNAs. Our Dataset comprises all 12 of these RNAs. Table 2 shows the type, sequence length and number of sequences in the MSA generated for each RNA target. Notably, synthetic RNAs in this dataset are longer, have more complicated folds, and exhibit fewer sequences in their MSAs than natural RNAs. Because of this, they present more difficult prediction challenges.

**Table 2.**
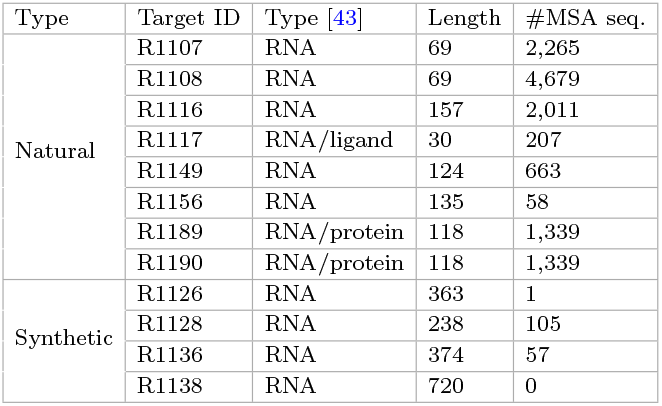
Properties of CASP15 RNA targets.

**Dataset 3**. Our aim was to create a dataset that could effectively evaluate the generalization abilities of the models. Since most tools do not explicitly disclose their training datasets, we used 13 January 2023, the validation set cutoff date for AlphaFold 3, as the training cutoff for all tools. This assumption is reasonable, as five other models were developed for CASP15 in April 2022, indicating they were trained on data published well before our chosen cutoff date. To compile this dataset, we selected structures published in the PDB database after 13 January 2023, ensuring that these RNAs were unseen during training. This approach enables a robust assessment of the models’ generalization ability to novel structures.

Starting with a download of all available RNA chains from PDB, which was 20, 320 RNA chains, we applied sequence identity clustering using MMseqs2 [44, 45], filtering for a minimum sequence identity of 90% and coverage of at least 80%, resulting in 3, 822 clusters. Clusters containing only RNAs published after 13 January 2023 were retained, followed by further filtering to remove sequences shorter than 16 nucleotides, with resolution greater than 9 Å, those with fewer than 90% defined residues, and those consisting solely of unknown residues (‘N’ or ‘X’). After these steps, 143 clusters remained, each uniquely represented by sequences with optimal resolution and maximal percentage of nucleotides with defined locations. This process is visualized in Figure 1.

**Fig. 1.**
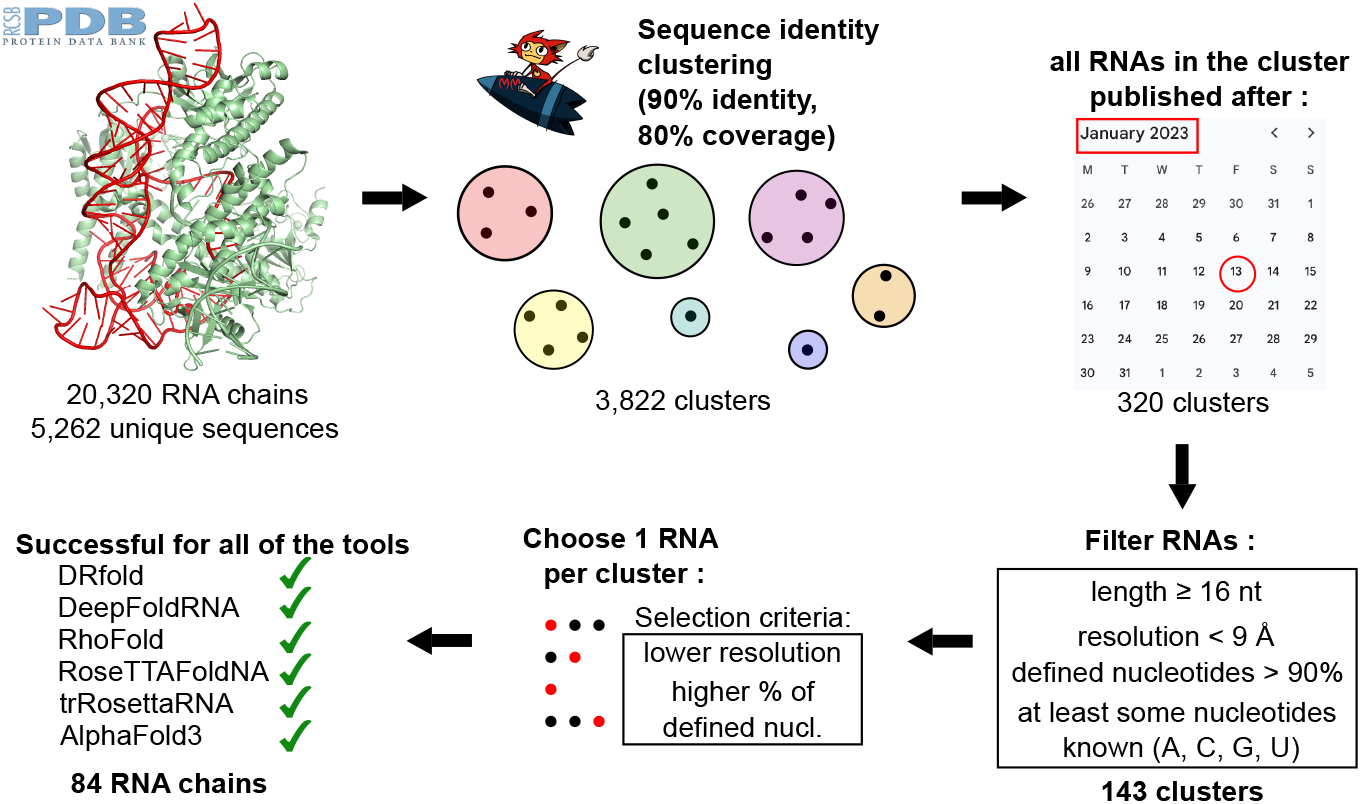
Process of creating Dataset 3. The process begins with extracting RNA chains from all complexes available in the Protein Data Bank (PDB). Next, the chains are clustered using MMseqs2 based on a sequence identity threshold of 90% and a minimum coverage of 80%. Clusters are then filtered to include only those where all sequences were published after January 13, 2023, ensuring that the dataset contains RNAs distinct from those used in the training datasets. Furthermore, several filtering criteria are applied to RNA chains and only those that satisfy all criteria are kept. From the remaining clusters, one representative RNA is selected per cluster, and predictions are attempted for all six tools. The final dataset consists of 84 RNAs for which all six tools successfully generated predictions.

For comparability, we only included sequences for which all models could produce final structures, yielding a final subset of 84 sequences. Notably, 15.5% of RNAs from the final dataset lacked MSA sequence coverage, and 63.1% had fewer than 100 MSA sequences, presenting more challenging tasks.

**Dataset 4** is a subset of Dataset 3, designed to include RNAs that are not only sequentially distinct from those in the training datasets but also structurally distinct. To identify these RNAs, we employed the RNA3DB pipeline [46], which clusters RNA structures based on both sequence and structural similarity.

The RNA3DB pipeline comprises four main steps: parsing, filtering, clustering, and splitting the data into training and test sets. Since our goal was to evaluate pre-trained models, we did not perform the splitting step and instead focused on using the structural clusters. We use the same first step, parse all RNA chains from PDB. To align the pipeline with our specific filtering criteria, we modified the filtering step by adjusting the minimum RNA length to 16 nucleotides and including NMR solutions that were not originally included.

During the clustering step, RNAs are first grouped based on 99% sequence identity using MMseqs2. Each representative sequence is then mapped to an Rfam family using Infernal with an E-value cutoff of 1.0, to capture structural homology. The resulting clusters, called components, are formed using the maximally connected subgraphs algorithm. If a representative RNA cannot be mapped to any Rfam family, it is assigned to a special component 0, which is typically recommended as a test set because these RNAs are structurally unique and thus suitable for assessing a model’s generalization ability.

After obtaining the clusters, we evaluated if each RNA from Dataset 3 was suitable for inclusion in Dataset 4. Specifically, we checked whether the RNA belonged to a cluster in which all member RNAs were published after 13 Jan 2023. This criterion ensured that none of the RNA’s structural relatives were part of the training datasets. Since Dataset 3 already enforced a sequence identity threshold of less than 90%, we retained the RNA3DB pipeline’s default 99% sequence identity threshold. This decision was made to avoid introducing false or missing edges in the graph that is built during the clustering process, as RNA3DB only checks structural similarity for the cluster representative. Most RNAs from Dataset 3 were assigned to component 0, reflecting their structural dissimilarity from other RNAs. Specifically, 58 out of 84 RNAs in Dataset 3 fell into component 0.

Dataset 4 comprises 60 RNAs in total, representing a highly stringent benchmark of structurally and sequentially distinct RNAs.

### 2.3 Metrics

While RMSD and TM-score [47] are widely recognized metrics, metrics like clash score [48] and local Distance Difference Test (lDDT) [49], and especially RNA-specific metrics such as Interaction Network Fidelity (INF) [50], add precision to our evaluation. These metrics offer supplementary insights into specific aspects of RNA structures, offering a well-rounded assessment.

To ensure accurate comparisons, we standardized both predicted and reference structures using RNA normalizer from RNA-Puzzles Toolkit [51].

**RMSD** (Root Mean Square Deviation) is a fundamental and widely employed metric for comparing two structures. It calculates the average distance between corresponding atoms in aligned predicted and reference structures. Here, we computed all-atom RMSD using the RNA-Puzzles Toolkit. RMSD values are reported in angstroms (Å), where 1 Å equals 10^−10^ meters, and lower scores indicate closer structural resemblance. RMSD values below 2 Å usually suggest high accuracy, while values between 2 −5 Å are commonly acceptable for broader structural assessments.

**TM-score** (Template-modeling score) serves as a metric for assessing the overall similarity between two structures, offering a global perspective compared to the localized sensitivity of RMSD. It emphasizes smaller structural differences more than larger ones, making it ideal for capturing global structural alignment. TM-score values range from 0 to 1 where a score of 1 indicates a perfect structural match, while scores above 0.45 denote similar folds, and values below 0.2 suggest a minimal similarity. We utilized USalign [52] to compute the TM-score, employing the alignment option based on the residue index from the PDB files, a crucial step for handling gaps in reference structures without misalignment.

**INF** (Interaction Network Fidelity) evaluates the fidelity of predicted interactions relative to the interactions from the reference structure, an important factor in RNA structure where Watson-Crick, non-Watson-Crick, and stacking interactions play key roles. INF values range from 0 to 1, with 1 indicating all interactions in the reference structure are correctly predicted, and values close to 0 suggesting significant deviations. We calculated INF using the RNA Puzzles Toolkit, in which essential base interactions are detected by MC-Annotate [53]. Different variants of INF (e.g., INF WC for Watson-Crick, INF NWC for non-Watson-Crick, and INF STACK for stacking interactions) allow us to assess models’ performance in predicting specific types of RNA interactions. This specificity is essential, as RNA structural stability relies heavily on accurately capturing these interactions.

The **clash score**, computed with MolProbity [48], quantifies atomic overlaps by calculating the number of clashing atoms per 1,000 atoms with overlaps greater than 0.4 Å. Lower clash scores indicate fewer steric clashes and greater physical plausibility of the structure. While a low clash score supports structural accuracy, it does not guarantee that the model captures the native structure perfectly; instead, it helps ensure the predicted conformation is free from unphysical atomic overlaps.

Finally, **lDDT** (local Distance Difference Test) scores were calculated using OpenStructure [54] to evaluate how well local atomic environments in the model match those in the reference structure. lDDT measures the preservation of atomic distances within a given inclusion radius, capturing local structural fidelity over distances up to 4 Å. The lDDT score ranges from 0 to 1, where values closer to 1 represent higher agreement with the reference structure’s local environment. This metric offers a residue-level assessment of structural accuracy, complementing the global orientation provided by RMSD and TM-score by highlighting the consistency of local atomic neighborhoods.

These metrics collectively provide a comprehensive view of RNA structure prediction quality, with each metric highlighting different structural characteristics and supporting a nuanced interpretation of each model’s performance.

### 2.4 Experiments

#### RNA 3D structure prediction

Most of the tools evaluated in this study predict only single-chain RNAs, making this our primary focus. Since none of the models have published their training code and training them is resource- and time-intensive, we relied on the published weights. We also used the default parameters for all tools. When a model outputs multiple structures, we select the first structure for comparison. Generating multiple sequence alignments (MSAs) can be time-consuming, so we generated the MSAs once using rMSA [31] and used them as input for all locally installed tools requiring MSAs, including DeepFoldRNA, RhoFold, RoseTTAFoldNA, trRosettaRNA and AlphaFold 3. For tools that use secondary structure as input, we followed their original pipelines without modifying the default secondary structure prediction model. When multiple structures are available in the PDB for the same RNA, we compare each predicted structure against all available references. For each prediction, we select the reference structure with the lowest all-atom RMSD and use this reference to report all other evaluation metrics. Additionally, we compared the performances of all of the tools on different length ranges and compared performances on single-chain RNAs and RNAs extracted from complexes.

#### Scoring functions

We aimed to determine whether the two most commonly used scoring functions, ARES and Rosetta score, could reliably identify the best prediction among the six. ARES is a deep learning-based scoring model for RNA structures, trained on 1000 FARFAR2-predicted structures [55] and their respective RMSDs across 18 RNAs. ARES is trained on RMSD values, meaning lower scores indicate better structural predictions. On the other hand, Rosetta score is an energy-based function from the Rosetta suite, designed to find the most energetically stable structure. Lower values are preferred here as well, although these scores are typically negative.

For our analysis, we first examined how well the scores from ARES and Rosetta correlate with RMSD. We also calculated the percentage of RNAs for which each scoring function selected the truly best prediction - the one with the lowest RMSD. Secondly, we incorporated ARES and Rosetta scores as additional “prediction tools” alongside the six RNA structure prediction models. For each RNA, we selected the prediction that received the best score from either ARES or Rosetta, and used the corresponding RMSD or TM-score as the data point in our visualizations. This approach allowed us to evaluate if the scoring functions could improve the overall structure prediction, even though they did not directly generate RNA structure predictions.

#### Complex prediction

Finally, since many RNAs from our Dataset 3 are extracted from complexes, we wanted to check at least with AlphaFold 3 and RoseTTAFoldNA, the only two tools that can predict complex structure, if the RNA chain predicted as part of the structure is better compared to when predicted as a single chain, without any context. Due to the time and resource demands of complex predictions, we used the AlphaFold 3 web server instead of the local version. The server imposes a limit of 5, 000 tokens, meaning nucleotides and amino acids combined. Consequently, some RNA chains extracted from large complexes, exceeding 5, 000 residues, could not be predicted as a full complex. In the end, there were 37 complexes, which covered 43 RNA chains from Dataset 3 that met these criteria for the AlphaFold 3 web server. On the other hand, while no strict residue limit is explicitly stated for RoseTTAFoldNA, generating a structure with over 1, 000 residues takes a couple of days. Additionally, predictions for complexes with over 2, 000 residues often appeared almost random, as shown for one example in Supplementary Figure S1. Based on these observations, we treated this as a restriction and limited ourselves to analyzing only complexes with up to 2, 000 residues. This resulted in 17 complexes and 19 RNA chains from Dataset 3.

## 3 Results and Discussion

### 3.1 RNA 3D structure prediction

#### Overall performance across the datasets

Figure 2 presents all-atom RMSD and TM-score metrics across Datasets 1, 2, and 3. In Figures 2A and 2B, evaluated on Dataset 1, which comprises RNA Puzzles, the models show their best performances, with low RMSD values and relatively high TM-scores. Overall, the tools perform similarly, with trRosettaRNA achieving the lowest mean RMSD of 6 Å, while RoseTTAFoldNA has the lowest median RMSD of 2.65 Å and the highest mean and median TM-score (0.601 and 0.688). These results suggest that Dataset 1 is the easiest for models to predict structures accurately, likely because it contains examples that may have been part of the training data. Interestingly, AlphaFold 3 lags significantly behind the other tools, with the highest median RMSD of 7.92 Å and the second-lowest median TM-score of 0.451. For more detailed visualization of all-atom RMSD and TM-score per RNA puzzle, refer to Supplementary Figure S2.

**Fig. 2.**
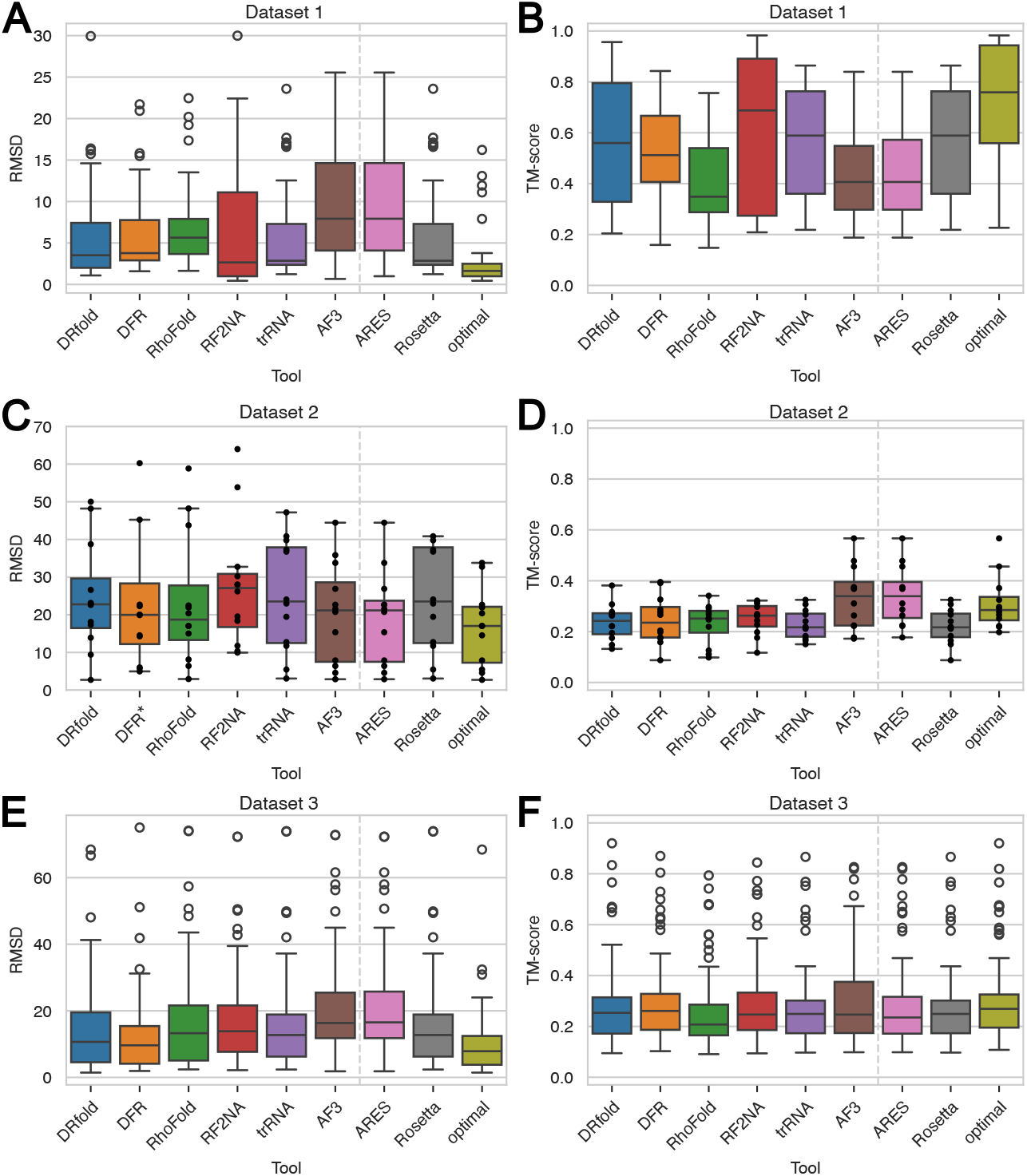
All-atom RMSD and TM-score for evaluated RNAs. Boxplot representation of all-atom RMSDs and TM-scores for RNA structure predictions by six deep learning-based methods. ARES and Rosetta refer to scoring functions, and for each RNA, the structure presented is the one with the best score among the six predictions according to these functions. The “optimal” column represents a hypothetical perfect scoring approach, where for each RNA the structure with the minimal RMSD among the six predictions is selected. Note that the corresponding TM-score shown for “optimal” selections is not necessarily the highest, as structures were chosen solely based on RMSD. A. All-atom RMSD for Dataset 1 (RNA Puzzles). B. TM-score for Dataset 1 (RNA Puzzles). C. All-atom RMSD for Dataset 2 (CASP15 RNA targets). *DeepFoldRNA structure for R1138, which had an RMSD of 178.6 Å, was excluded from the visualization. D. TM-score for Dataset 2 (CASP15 RNA targets). E. All-atom RMSD for Dataset 3. F. TM-score for Dataset 3. *Abbreviations used: DFR = DeepFoldRNA, RF2NA = RoseTTAFoldNA, trRNA = trRosettaRNA, AF3 = AlphaFold 3*.

In contrast, Figures 2C and 2D for Dataset 2, CASP15 RNA targets, reveal the poorest performances, with TM-scores approaching values seen in random folds. The high RMSD values, including an extreme case for R1138 (DeepFold’s model has RMSD = 178.6 Å), underscore the complexity of Dataset 2 and raise questions about the models’ capacity to model complex and large molecules. Supplementary Figure S3 provides additional detail, showing all-atom RMSD and TM-scores for individual RNA targets in Dataset 2. Notably, synthetic RNAs, which appear as outliers in Figure 2C, have the worst predicted structures. For synthetic RNAs, the median RMSD ranges from 34.81 Å for AlphaFold 3 to 52.74 Å for DeepFoldRNA, whereas for natural RNAs, RMSDs are lower, from 11.62 Å (AlphaFold 3) to 17.7 Å (DRfold). The discrepancy likely reflects the more common appearance of natural RNAs in the PDB and, consequently, in the training datasets, making synthetic RNAs more challenging for these models. Interestingly, AlphaFold 3 performs relatively well on synthetic RNAs, suggesting a potential advantage in certain unseen cases. However, given that Dataset 2 contains only 12 examples, its small size limits definitive conclusions about the models’ broader generalization abilities.

Finally, we specifically curated Dataset 3 to assess the models’ generalization abilities. Figure 2E shows that all-atom RMSD values are centered around a median of 13 Å, ranging from 9.64 Å for DeepFoldRNA to 16.33 Å for AlphaFold 3, indicating moderate prediction accuracy but still leaving room for improvement. On the other hand, Figure 2F shows generally low TM-scores across the tools, with median values spanning between 0.207 for RhoFold to 0.261 for DeepFoldRNA, with only a single prediction by DRfold among all achieving TM-score above 0.9. These results underscore the difficulty in achieving reliable generalization beyond training data for these tools. An example of predictions for one RNA from Dataset 3 is shown in Figure 3.

**Fig. 3.**
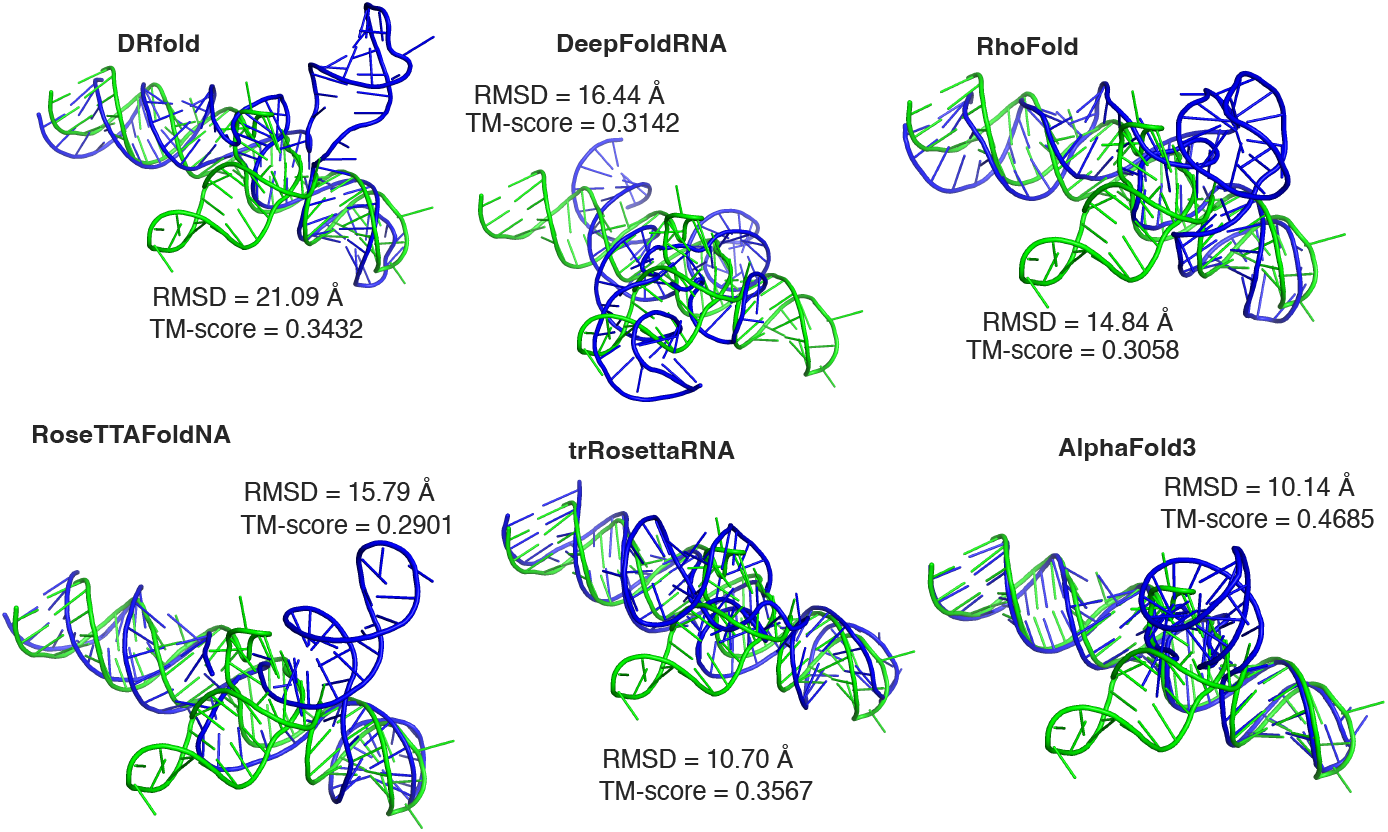
Predictions for RNA chain (PDB_ID: 8T29_R). The native structure is shown in green and the predicted structures in blue. All six predicted structures for this RNA chain exhibit relatively high RMSD and low TM-scores. Among these, AlphaFold 3 produced the most accurate prediction, capturing a reasonable global fold. ARES selected AlphaFold 3’s prediction as the best, while Rosetta score favored trRosettaRNA’s structure. For this RNA, AlphaFold 3’s prediction indeed had the lowest all-atom RMSD and the highest TM-score, but this is often not the case. Visualizations of the structures were generated using PyMOL [56].

#### Performance based on other important metrics

It is important to note that while RMSD and TM-score offer some insight into structural accuracy, they can yield quite different results for the same predictions. To gain a more comprehensive understanding, we further evaluated these predictions using a range of metrics, including RNA-specific ones. Supplementary Figure S4 illustrates the correlations between various metrics for Dataset 3. For consistency in correlation direction, we inverted the values of RMSD and clash score, as lower values are preferred for these metrics. Interestingly, the only high correlation is between INF ALL and INF STACK. This is likely because stacking interactions are present in all RNAs in the dataset, making INF STACK most influential on INF ALL. The lack of high correlations among the metrics suggests that each metric captures unique structural information, highlighting the value of using multiple metrics to evaluate predictions. This also implies that different tools may perform best depending on the specific structural aspect under consideration.

Figure 4 illustrates additional evaluation metrics for Dataset 3, including INF ALL, INF WC, INF NWC, INF STACK, clash score, and lDDT. Results using the same metrics for Datasets 1 and 2 are available in Supplementary Figures S5 and S6. When presenting the INF metrics, the RNA Puzzles toolkit assigns a value of −1 when neither the reference nor the predicted model contains a specific type of interaction. To ensure clarity and consistency in visualization, we excluded RNAs lacking certain interaction types. Additionally, if the reference model includes an interaction that the predicted model fails to capture, we assigned an INF value of 0, treating these instances as significant prediction failures.

**Fig. 4.**
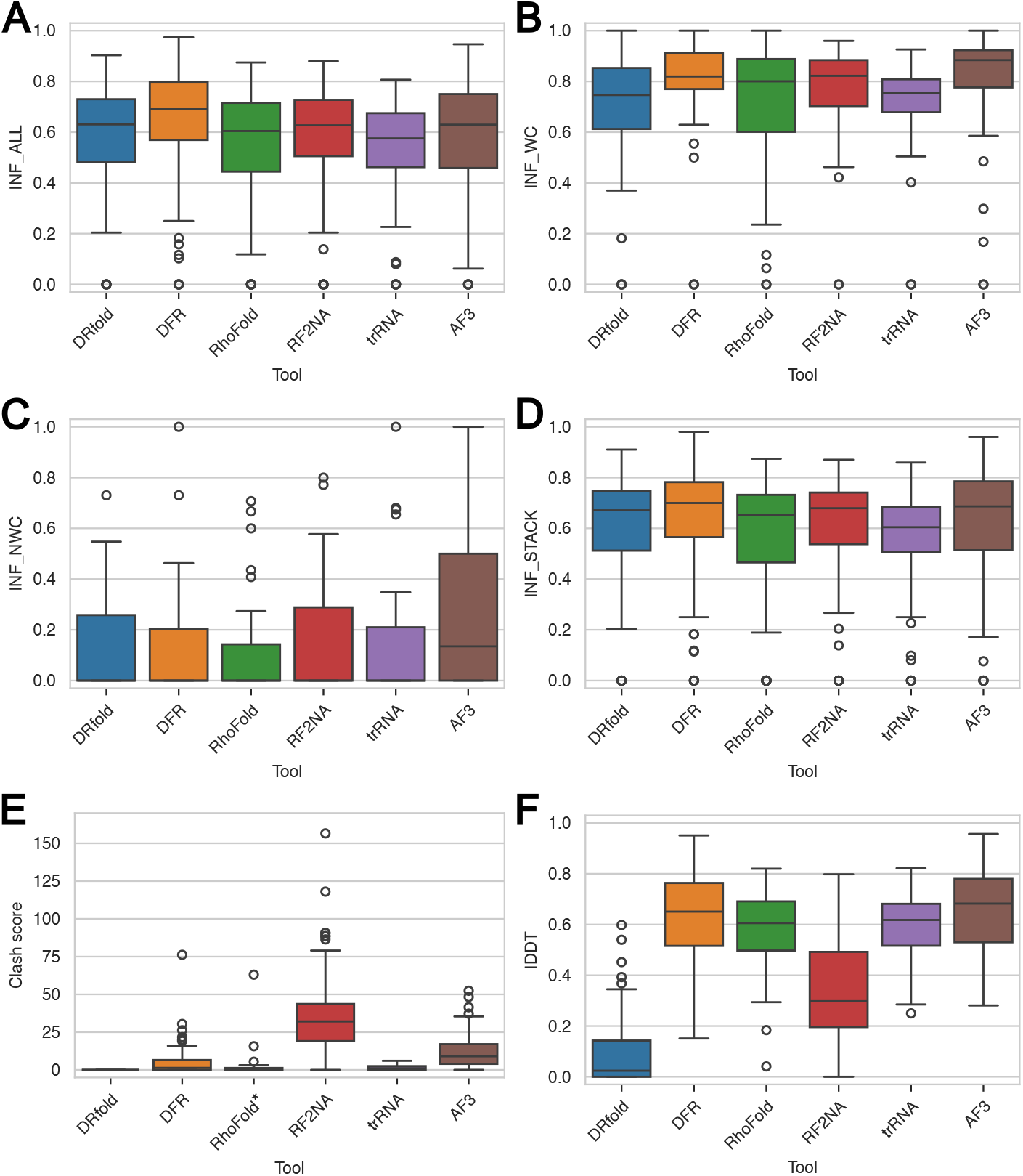
Other important metrics for Dataset 3. For INF metrics, all RNAs that do not have a certain type of interaction are excluded from visualizations. A. INF ALL (all RNAs have some interactions). B. INF WC (excludes 47 RNAs without any Watson-Crick interactions). C. INF NWC (excludes 55 RNAs without any non-Watson-Crick interactions). D. INF STACK (all RNAs have stacking interactions). E. Clash score. *RhoFold structures with very large clash scores were excluded from the visualization. F. lDDT. *Abbreviations used: DFR = DeepFoldRNA, RF2NA = RoseTTAFoldNA, trRNA = trRosettaRNA, AF3 = AlphaFold 3*.

For Dataset 3, all tools perform best for Watson-Crick interactions (see Figure 4B) and worst for non-Watson-Crick interactions (see Figure 4C). Median values for all interactions range from 0.575 for trRosettaRNA to 0.691 for DeepFoldRNA. DeepFoldRNA also performs the best for stacking interactions with a median of 0.700. AlphaFold 3 outperforms other tools by a large margin for non-Watson-Crick interactions, with a median value of 0.297 compared to 0.119 to 0.145 for the rest of the tools. AlphaFold 3 is also the best for Watson-Crick interactions, with a very high median of 0.884. In Figure 4E, an outlier for RhoFold with a clash score of 196.93 was removed from visualizations. While lower clash scores generally indicate better structural accuracy, Drfold consistently reports a clash score of 0, even though some native structures have a clash score larger than 0. For instance, more than 83% of references in Dataset 3 have clash scores greater than 0. RoseTTAFoldNA has the highest median clash score (32.09), and AlphaFold 3 second-highest (9.06). In terms of the lDDT metric, shown in Figure 4F, DRfold performs the worst (median lDDT = 0.024), whereas AlphaFold 3 exhibits the best performance among the evaluated tools (median lDDT = 0.683).

Overall, DeepFoldRNA demonstrates the strongest performance across the majority of metrics for Dataset 3. Specifically, it achieves the best results for all-atom RMSD, TM-score, INF all, and INF STACK, while also maintaining a respectable clash score (third best, with a median of 0.76) and good lDDT, where it is second best with a median of 0.651. The only metrics where it falls short are INF WC and INF NWC, where AlphaFold 3 and RoseTTAFoldNA outperform it.

Supplementary Figure S5 for Dataset 1 reveals that RoseTTAFoldNA achieves the best results for INF ALL, INF STACK and INF NWC. However, DeepFoldRNA slightly outperforms RoseTTAFoldNA in INF WC. For lDDT, AlphaFold 3 achieves the highest score, while RoseTTAFoldNA stands out with a relatively high median clash score. Metrics for Dataset 2 are illustrated in Supplementary Figure S6, where AlphaFold 3 demonstrates superiority across all evaluated metrics, except for clash score, where it has the second-highest value. Despite this exception, AlphaFold 3 performs the best overall for the CASP15 RNA targets.

#### Assessing generalization abilities on Dataset 4

To highlight the impact of splitting datasets based solely on sequence similarity versus on both structural and sequence similarity, we compare all-atom RMSD (in Figure 5A) and TM-score (in Figure 5B) across Dataset 3 and its subset, Dataset 4. While median all-atom RMSD values remain consistent between the two datasets, indicating similar overall trends, TM-scores on Dataset 4 show a slight decrease, with DeepFoldRNA emerging as the top performer, achieving a median of 0.231. This suggests that factoring in both structural and sequence similarity when creating subsets may be important for accurately assessing model performance, as it can reveal subtler distinctions in prediction quality that a sequence-only split might overlook. Other metrics for Dataset 4 are shown in Supplementary Figure S7. Given that Dataset 4 comprises over 70% of Dataset 3, it is unsurprising that the overall conclusions remain consistent. DeepFoldRNA continues to outperform other models across most metrics, confirming it generalizes better than the other models. However, the results still fall short of desirable levels, with median RMSD values around 10 Å and median TM-scores only slightly exceeding 0.2, which are comparable to random fold values.

**Fig. 5.**
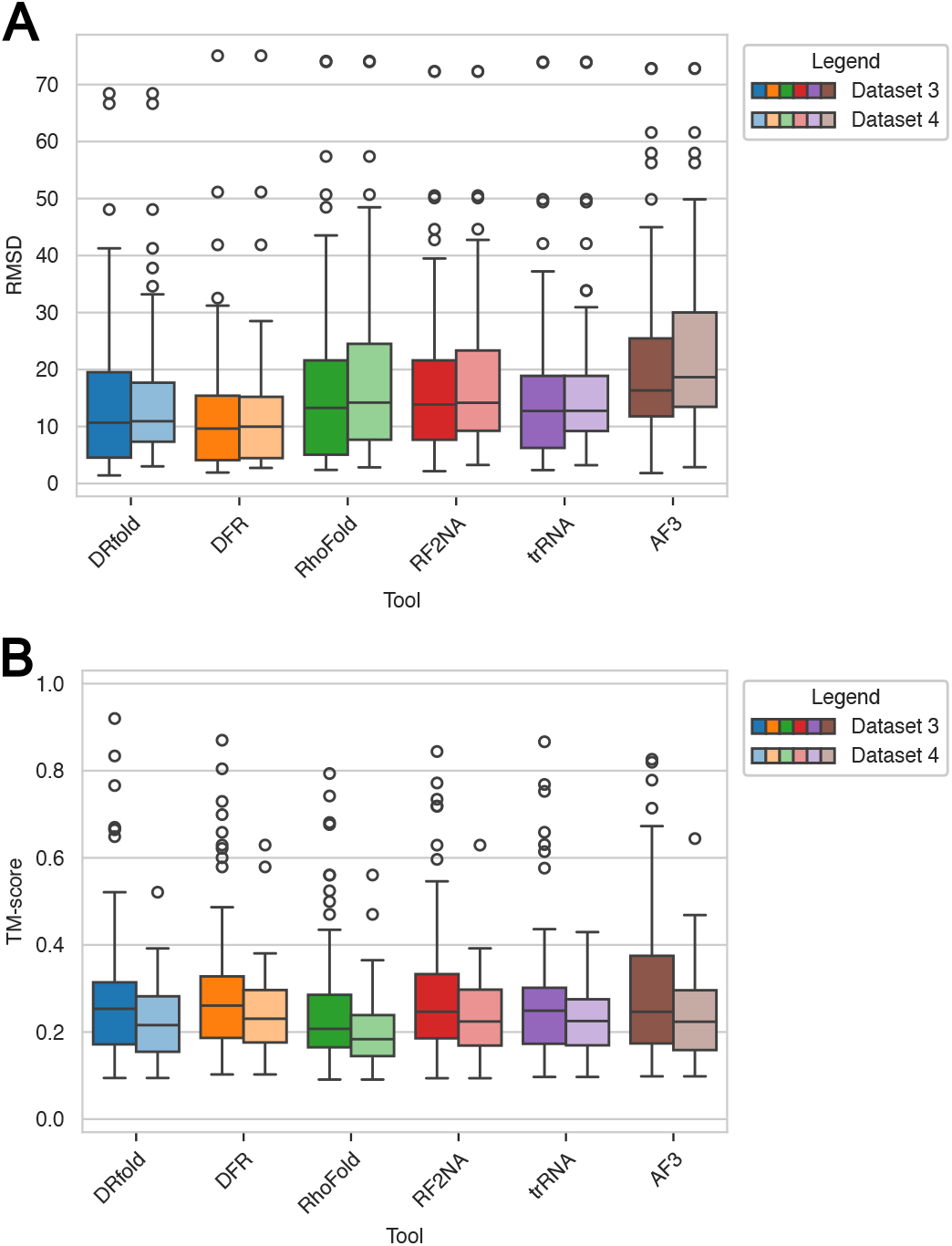
Comparison of all-atom RMSD and TM-score for Dataset 3 and its more stringent subset Dataset 4. There are 84 RNA chains in Dataset 3 and 60 RNA chains in Dataset 4. A. RMSD for Datasets 3 and 4. B. TM-score for Datasets 3 and 4. *Abbreviations used: DFR = DeepFoldRNA, RF2NA = RoseTTAFoldNA, trRNA = trRosettaRNA, AF3 = AlphaFold 3*.

While some INF metrics, such as INF WC, show promising median values above 0.84, there remains significant room for improvement, particularly in INF NWC. Future efforts should focus on enhancing the models’ ability to accurately predict stacking interactions and, most critically, complex non-Watson-Crick interactions, which are essential for capturing biologically accurate RNA conformations.

#### Performance per different RNA length groups

To assess the impact of RNA length on prediction accuracy, we divided RNAs from Dataset 3 into length-based groups: 16 −30, 30 −50, 50 −80, and 80 −135 nucleotides. Figure 6 presents all-atom RMSD (Figure 6A) and TM-score (Figure 6B) for each group. The best results are observed for the shortest group, consisting of 16 to 30 nucleotides, and the worst overall results are in the longest group, between 80 and 135 nucleotides, but the median values did not increase significantly with each successive length group. This could be because even the longest RNA in the dataset is not classified as really long, exceeding 200 nucleotides. In each length group, DeepFoldRNA is among the first two best tools.

**Fig. 6.**
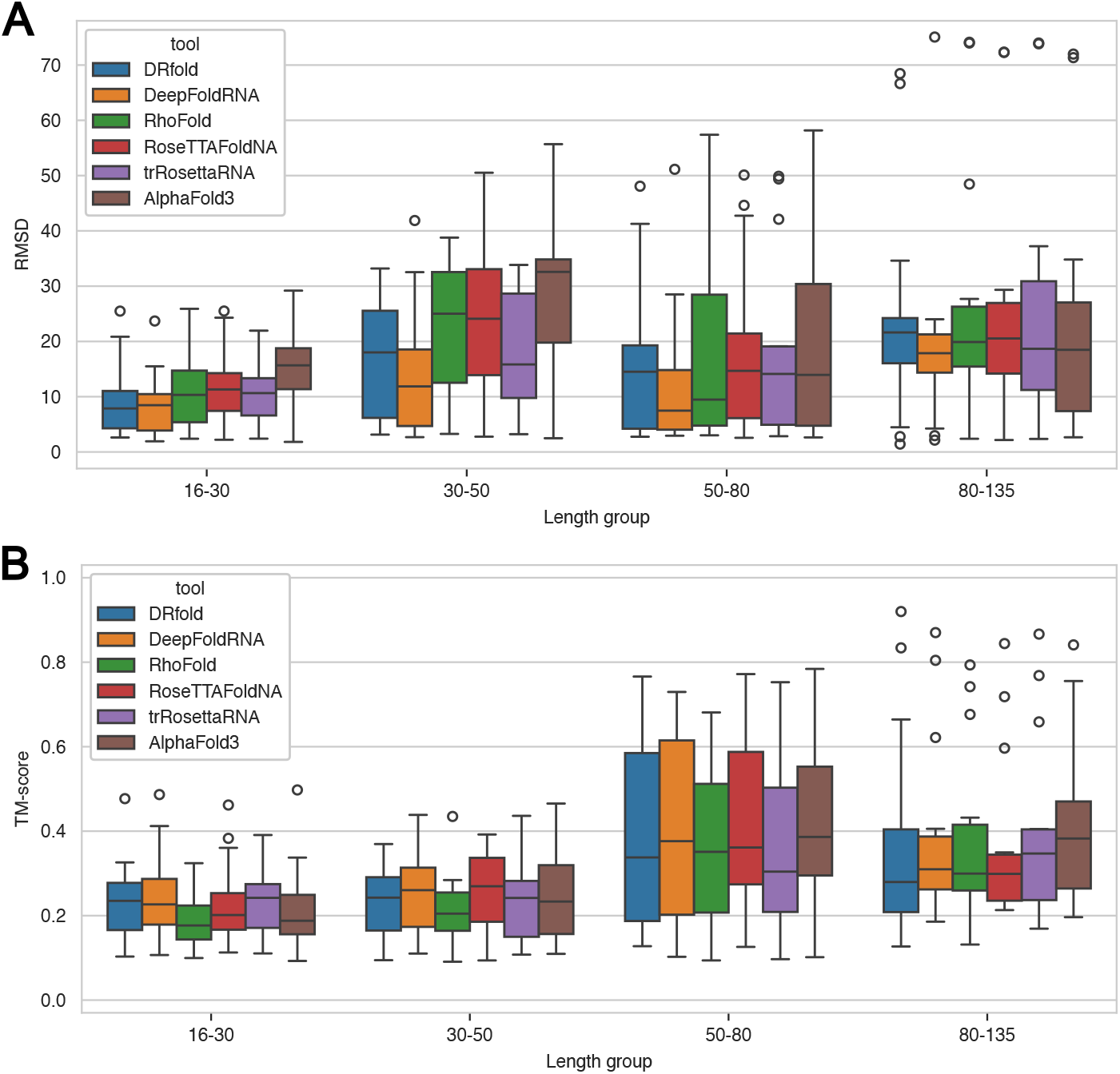
RMSD and TM-score for Dataset 3 per length group. There are 39 RNAs with lengths between 16 and 30, 16 RNAs with lengths between 30 and 50, 15 RNAs with lengths between 50 and 80, and 14 RNAs with lengths between 80 and 135. A. All-atom RMSD. B. TM-score. *Abbreviations used: DFR = DeepFoldRNA, RF2NA = RoseTTAFoldNA, trRNA = trRosettaRNA, AF3 = AlphaFold 3*.

TM-scores are lowest for the shortest RNAs, in contrast to RMSD results. This observation aligns with findings from other studies, which suggest that the TM-score is more stringent for very short RNAs (length *<* 30) [57]. Interestingly, the longest RNAs include a few examples with the highest TM-scores. However, these three cases, two 5S ribosomal RNAs and one tRNA, are common in training datasets. While they appear sequentially distinct, they are not structurally unique compared to the training data and are excluded from Dataset 4. This highlights the importance of a structural split, as these outliers could lead to different conclusions if not properly accounted for.

#### Performance across different RNA types

To evaluate model performance across different RNA types, we examined single-chain RNAs and RNA chains from complexes, all from Dataset 3. Figure 7 presents the all-atom RMSD and TM-score results for each category. The dataset includes 11 single-chain RNAs, and 73 RNA chains from complexes, where only 2 come from RNA-RNA complexes, and the rest is from protein-RNA complexes.

**Fig. 7.**
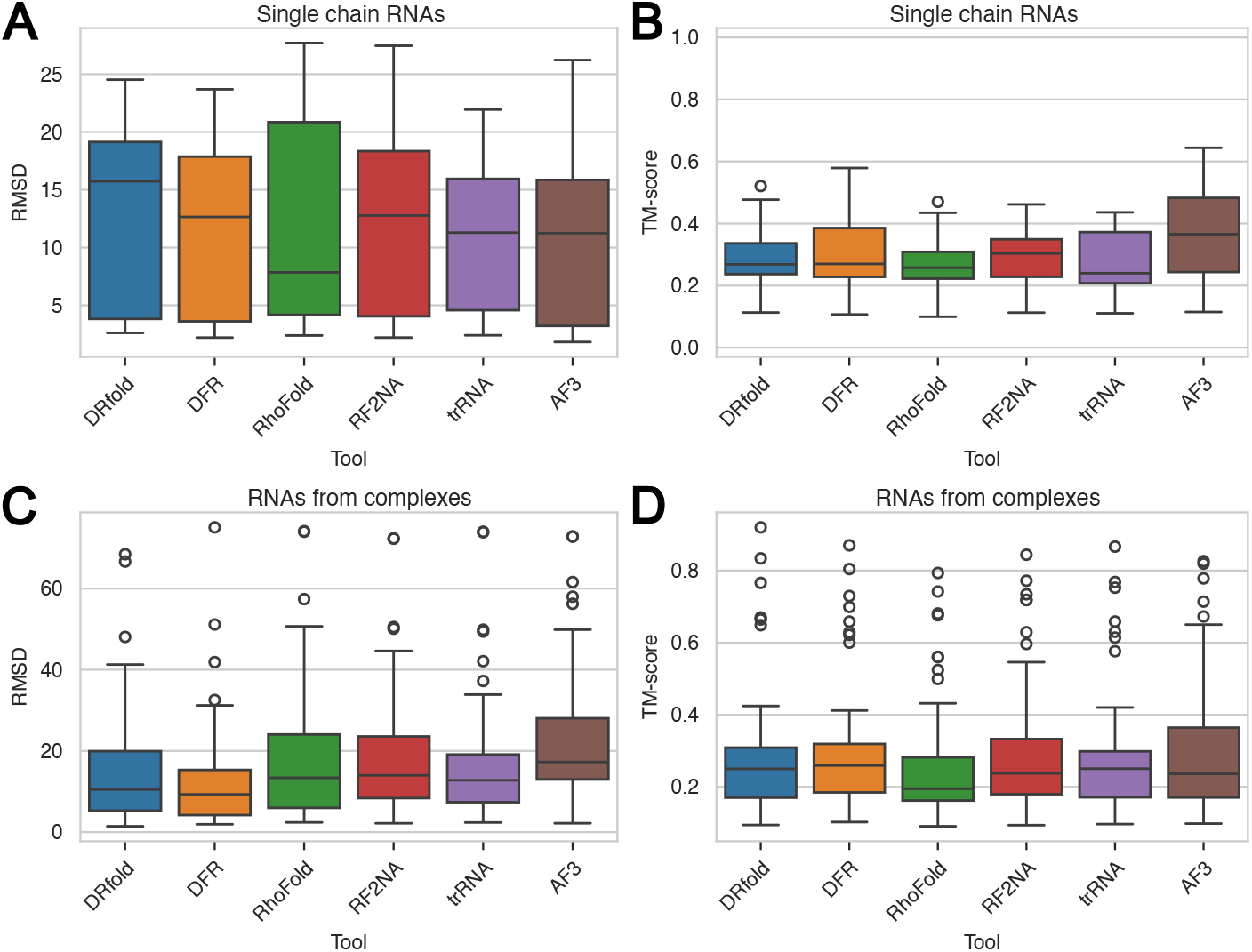
All-atom RMSD and TM-score for different RNA types: single chain RNAs and RNA chains extracted from complexes. A. RMSD for single chain RNAs from Dataset 3 (11 RNA chains). B. TM-score for single chain RNAs from Dataset 3 (11 RNA chains). C. RMSD for RNA chains from complexes from Dataset 3 (73 RNA chains). D. TM-score for RNA chains from complexes from Dataset 3 (73 RNA chains). *Abbreviations used: DFR = DeepFoldRNA, RF2NA = RoseTTAFoldNA, trRNA = trRosettaRNA, AF3 = AlphaFold 3*.

For single-chain RNAs, AlphaFold 3 stands out with the highest median TM-score of 0.366, significantly above the second-best, RoseTTAFoldNA with a median TM-score of 0.304, while RhoFold achieves the best median RMSD of 7.85 Å. In contrast, RNA chains from complexes generally yield lower-quality predictions; however, their median RMSD values are comparable to those of single-chain RNAs. Among the tools, DeepFoldRNA performs best for complex-derived RNAs, with a median RMSD of 9.44 Å and a TM-score of 0.259.

RNAs from complexes exhibit some examples with excellent TM-scores, a pattern not observed for singlechain RNAs. This difference could be attributed to the limited number of single-chain RNA examples (only 11) and the fact that certain complex-derived RNAs, such as 5S rRNA and tRNA, share similar global folds with training data. Although sequentially distinct, these RNAs have well-known global folds, which might explain better performance for most of the tools.

#### Performance overview and a way forward

Overall, it is difficult to definitively say that one tool outperforms the others, as different tools excel for different datasets and metrics. Most tools perform similarly, and their results still fall short of the accuracy achieved in protein structure prediction, leaving substantial room for improvement. Given their comparable performance, we investigated which tool performs best for each RNA from Dataset 3 in terms of RMSD and TM-score, with the results shown in Figure 8. The figure illustrates that all six tools predict the best structure for some RNAs, and same as before, the results vary depending on the evaluation metric used. For instance, Figure 8B shows that while AlphaFold 3 often achieves the highest TM-scores, its predictions do not consistently perform as well when evaluated by all-atom RMSD. The same analysis for Dataset 1 and Dataset 2 are shown in Supplementary Figures S8 and S9. These findings prompted us to explore the possibility of selecting the best prediction by combining tools and scoring the outcomes, potentially leading to higher-quality predictions.

**Fig. 8.**
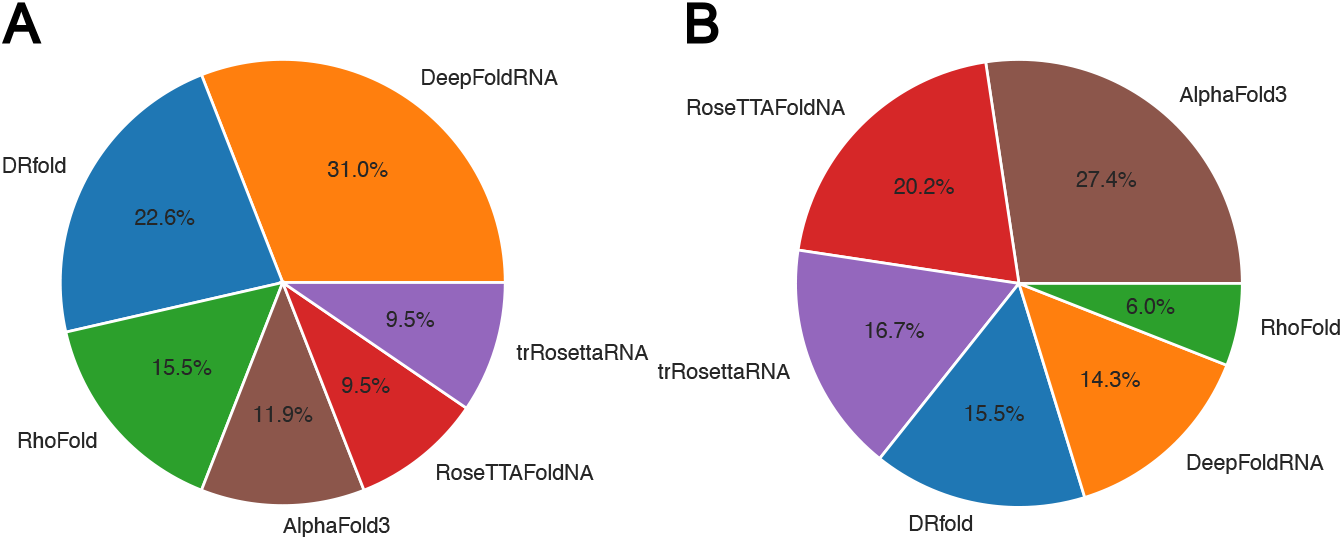
Percentage of RNA chains for which each tool predicted the best structures according to lowest RMSD and highest TM-score for Dataset 3. A. The best structure for each RNA is determined according to all-atom RMSD. B. The best structure for each RNA is determined according to TM-score.

### 3.2 Scoring functions

For scoring, we used ARES and Rosetta scores. In Figure 2, each panel (A–E) includes a gray dashed line, followed by boxes that represent RMSD and TM-score for the predictions scored as the best by these two scorers. These are followed by a box labeled “optimal”, representing a hypothetical perfect scoring approach that selects the structure with the lowest RMSD among the six predictions for each RNA. This inclusion illustrates the potential improvement achievable with an optimal scoring method and highlights the best possible results attainable with the current prediction models.

Rosetta score predominantly selects trRosettaRNA’s predictions, likely because trRosettaRNA’s refinements are guided by Rosetta’s own energy function. As a result, the Rosetta scoring column looks the same as trRosettaRNA’s results in Datasets 1 and 3. For Dataset 2, Rosetta score chose DeepFoldRNA’s prediction for R1138 as the best, despite this structure’s exceptionally high RMSD of 178.6 Å, leading Rosetta score to select the best structure in only 10.45% of RNAs overall (13.51% for Dataset 1, 8.33% for Dataset 2 and 9.52% for Dataset 3). Spearman correlation between Rosetta score and RMSD is −0.0116 for Dataset 1, 0.1810 for Dataset 2 and −0.0027 for Dataset 3, with these low values indicating that the predictions of structural accuracy are similar to a random choice.

ARES, commonly used by many groups in CASP competitions, was somewhat better. We noticed that it favored AlphaFold 3’s predictions in many cases, or to be precise, for all but one RNAs in Dataset 2, all but two RNAs in Dataset 1, and 83.33% RNAs in Dataset 3. Thus, it selected the best structure only for only 16.48% of all RNAs (5.41% of Dataset 1, 33.33% in Dataset 2 and 10.71% in Dataset 3). Spearman correlation between ARES scores and RMSD is 0.0147 for Dataset 1, 0.3571 for Dataset 2, and −0.2190 for Dataset 3. These poor results may come from ARES being trained exclusively on structures generated using FARFAR2, suggesting that retraining it with structures predicted by tools benchmarked here could enhance its performance.

The optimal scoring function would significantly reduce the average RMSD and improve TM-scores across datasets. For Dataset 1, the average RMSD would decrease from 6.01 Å, achieved by trRosettaRNA, to 3.01 Å, while the average TM-score would increase from 0.601, achieved by RoseTTAFoldNA, to 0.721. On Dataset 2, the average RMSD would improve from 20.255 Å for AlphaFold 3 to 16.730 Å. However, discrepancies between metrics become evident when comparing the TM-scores for Dataset 2; the average TM-score for structures selected based on minimum RMSD would drop slightly to 0.312 compared to AlphaFold 3’s 0.332. For Dataset 3, an optimal scorer would reduce the average RMSD from 12.12 Å for DeepFoldRNA to 9.83 Å, and the TM-score would slightly improve from 0.296 (AlphaFold 3) to 0.300. These results highlight the potential for improvement with a perfect scoring function and the challenges posed by conflicting metrics.

We conclude that the existing scoring methods do not significantly aid in consistently selecting the best structure among model predictions, and we believe that working on such scoring functions could benefit the field.

### 3.3 Complex prediction

Most tools predict only single RNA chains, yet the majority of RNA chains in the PDB come from complexes. This led us to investigate whether predictions for these chains improve when the entire complex is predicted, as opposed to predicting the RNA chain alone. This comparison was possible only for AlphaFold 3 and RoseTTAFoldNA. As previously noted, AlphaFold 3 enabled us to predict 37 complexes, from which we extracted 43 RNA chains, while RoseTTAFoldNA allowed for predictions of 17 complexes, from which we extracted 19 RNA chains. Figure 9 compares the RMSD values for chains predicted within a complex versus as single RNA chains.

**Fig. 9.**
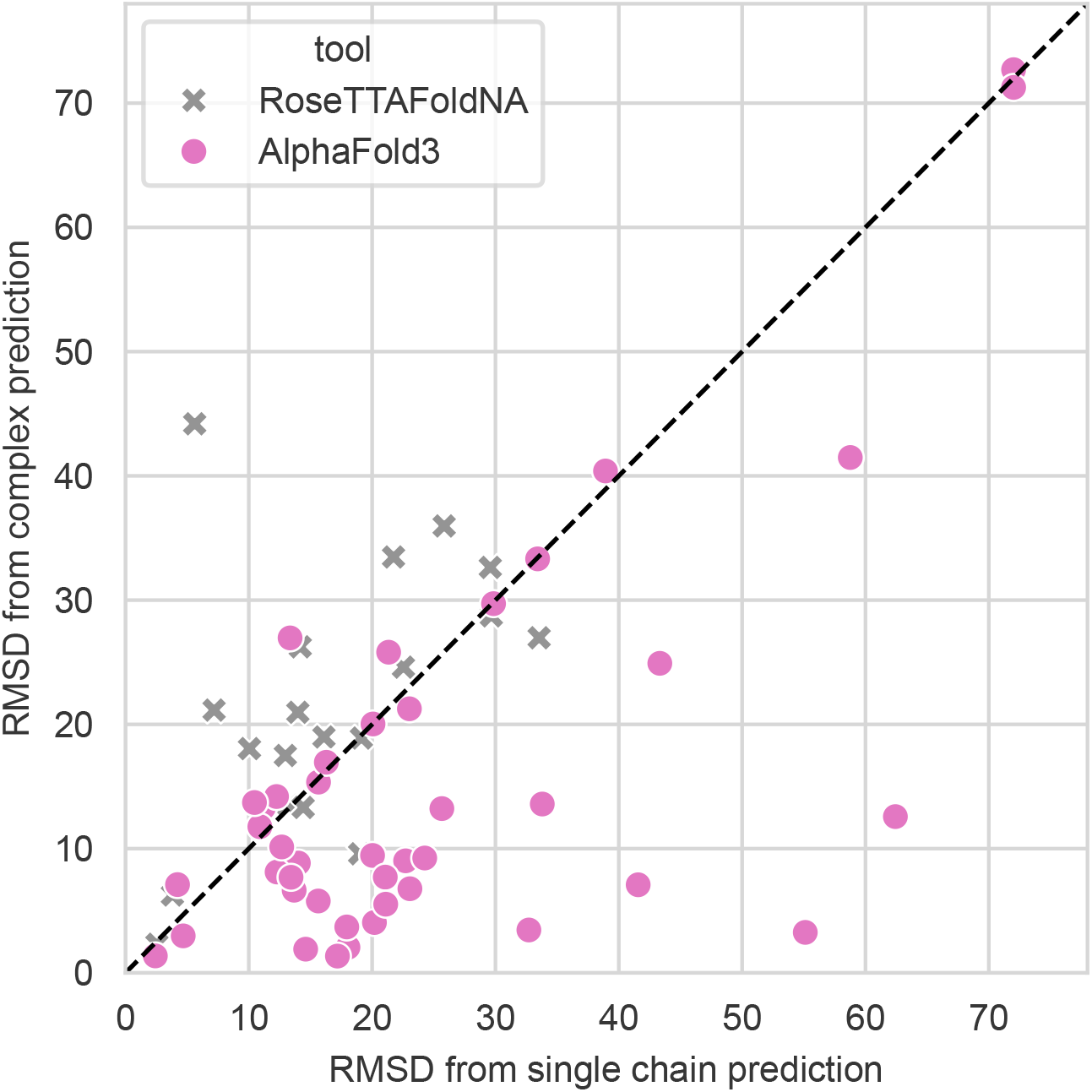
All-atom RMSD for structures of RNA chains predicted as a single chain and part of a complex. There are 43 RNA chains predicted by AlphaFold 3 and 19 RNA chains predicted by RoseTTAFoldNA. Data points below the diagonal line indicate that the prediction made as part of a complex has a lower RMSD and is thus more accurate, whereas points above the line signify that the single-chain prediction yields a better result.

For AlphaFold 3, predictions within the full complex are notably more accurate than those for isolated RNA chains. However, for RoseTTAFoldNA, this trend does not hold; in fact, single-chain predictions often yield better results than those from complex predictions. These unexpected differences may stem from the varying proportions of single-chain and complex examples in each model’s training dataset, even though both models were trained on both complex and single-chain RNA structures.

To provide a broader comparison of complex prediction results, we present an overview of both AlphaFold 3 and RoseTTAFoldNA performance in complex prediction mode in Figure 10. Figures 10A and 10B show the all-atom RMSD and TM-score, respectively, for the 43 RNA chains predicted using AlphaFold 3. Figures 10C and 10D present the all-atom RMSD and TM-score for the 19 RNA chains predicted using RoseTTAFoldNA in complex mode.

**Fig. 10.**
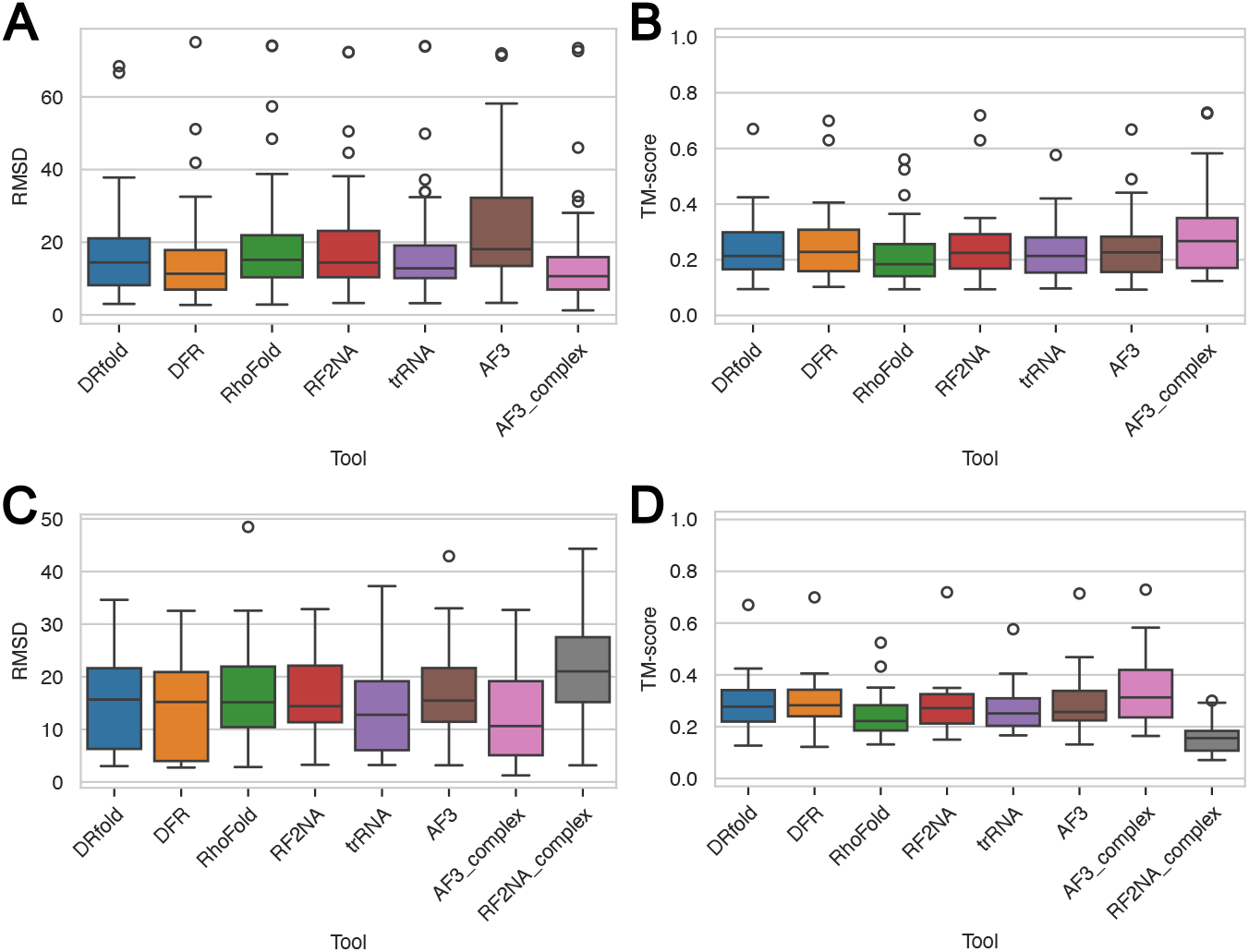
RMSD and TM-score for RNA chains from some complexes from Dataset 3. The suffix “complex” indicates predictions for the entire complex, not just the RNA chain within it. A. All-atom RMSD for 43 RNA chains from complexes that can be predicted by AlphaFold 3 (complexes with fewer than 5000 residues). B. TM-score for the same 43 RNA chains from complexes predicted by AlphaFold 3 (complexes with fewer than 5000 residues). C. All-atom RMSD for 19 RNA chains from complexes. D. TM-score for the same 19 RNA chains from complexes. *Abbreviations used: DFR = DeepFoldRNA, RF2NA = RoseTTAFoldNA, trRNA = trRosettaRNA, AF3 = AlphaFold 3*.

The results show that AlphaFold 3’s complex predictions yield the best outcomes, achieving superior accuracy in both RMSD and TM-score. Supplementary Figures S10 and S11 further confirm this, showing additional metrics for these subsets. Notably, AlphaFold 3 demonstrates strong performance in INF metrics, particularly for non-Watson-Crick interactions, highlighting its advantage in complex prediction accuracy.

### 3.4 Computation time

While performance is critical to assess, computational efficiency is equally important for practical applications, especially in scenarios where structure prediction occurs repeatedly within a training loop, as in models like gRNAde [58] for inverse folding. High execution times in such cases can pose significant challenges.

Since generating MSAs can be time-consuming, we opted to compute them once and use the same MSAs across all tools that require them: DeepFoldRNA, RhoFold, RoseTTAFoldNA, trRosettaRNA and AlphaFold 3.

This approach introduces some limitations in fair time comparisons, as DRfold does not require MSAs. Nonetheless, we believe that presenting an execution time overview is still informative. Supplementary Figure S12 shows execution times (in hours) for Dataset 3 RNAs for all six tools, as all were run locally.

Among the tools evaluated, DeepFoldRNA typically requires the longest runtime, with both DeepFoldRNA and DRfold exhibiting exponential increases in execution time as RNA length increases. For instance, processing the longest RNA in our dataset, comprising 135 nucleotides, took more than 7 hours for DeepFoldRNA and more than 3 hours for DRfold to process. In some cases not included in our datasets, DRfold required nearly 100 hours (over four days) to process RNAs slightly exceeding 300 nucleotides. For both tools, the time-intensive step is L-BFGS optimization: this step is essential for DeepFoldRNA, which only outputs geometrical restraints. In DRfold, however, frames are produced, but then DRfold combines these frames and a geometrically informed energy function that it then minimizes, adding substantial computation time.

While execution time is a critical factor in practical applications, its importance depends on the specific use case. Generating MSAs is particularly time-intensive, making pipelines that perform well with only single-sequence inputs highly desirable for tasks where rapid predictions are prioritized. For scenarios where accuracy is more critical than speed, the additional computational time required by methods like L-BFGS optimization or refinement may be justifiable. However, introducing flexibility into prediction pipelines, for example, an optional flag to toggle resource-intensive steps like MSA generation or optimization, could offer a balanced approach, allowing users to adapt the pipeline based on their specific requirements, whether for exploratory analysis or higher-precision predictions.

## 4 Conclusion

This study benchmarked six state-of-the-art deep learning-based tools for RNA 3D structure prediction across four diverse datasets, employing a range of metrics such as all-atom RMSD, TM-score, INF metrics, clash score, and lDDT. The evaluation revealed varied performance among tools, influenced by dataset composition, RNA characteristics, and the choice of metric, highlighting the inherent challenges in comprehensive comparisons.

Dataset-specific analysis showed that tool performance can vary significantly. On Dataset 1, which comprises the RNA Puzzles, most tools perform similarly well, with AlphaFold 3 slightly underperforming compared to others. The generally high performance on this dataset is likely due to overlaps between the RNA Puzzles and the tools’ training datasets. On Dataset 2, consisting of CASP15 RNA targets, AlphaFold 3 emerged as the top performer across most metrics, except for clash score. It excelled in predicting synthetic RNAs, but with RMSD values exceeding 30 Å and TM-scores below 0.5. AlphaFold 3 was also the best for natural RNAs within Dataset 2. Evaluations on the proposed Dataset 3 highlighted DeepFoldRNA as the overall best-performing tool, excelling across most metrics except INF WC and INF NWC, where AlphaFold 3 showed better results. Finally, Dataset 4, a subset of Dataset 3 emphasizing structural and sequence dissimilarity, confirmed DeepFoldRNA’s best generalization capabilities among the tools, with performance trends consistent with those observed in Dataset 3.

Tool performance also depended on RNA chain context and length. For shorter RNAs, predictions generally achieved better RMSD values, while TM-scores were highest for RNAs between 50 and 80 nucleotides, reflecting the metric’s stricter evaluation of very short RNAs. Since the longest RNA in our dataset is only 135 nucleotides, we cannot discuss what happens with prediction accuracy for really long RNAs.

While we anticipated the best performance for single-stranded RNAs, some RNAs from complexes yielded remarkably good results. This could be attributed to the structural similarity of these specific RNA examples, which were well-predicted and may have influenced the overall conclusions.

Further experiments comparing predictions for RNA chains modeled as single entities versus as part of a complex revealed that AlphaFold 3 performed significantly better when predicting full complexes. This suggests that incorporating structural context during model training could substantially improve predictions, especially for RNAs within complex environments.

Our evaluation of the ARES and Rosetta scoring functions showed limited success in reliably identifying the best predictions among the six tools. This underscores a critical area for improvement in RNA structure prediction: the development of robust scoring functions to aid in selecting the most accurate structures. Despite their limitations, each structure prediction tool demonstrated unique strengths, with all six occasionally producing the best predictions for specific RNAs. This diversity suggests that rather than striving to identify a universally superior tool, future efforts should focus on strategies to integrate or select the best predictions based on the specific context and available data.

Overall, while the evolution of deep learning-based tools for RNA structure prediction is promising, significant work remains to refine their performance and address existing limitations. Accurate RNA structure prediction is crucial not only for advancing our understanding of RNA biology but also for unlocking new possibilities in drug discovery, RNA-based therapeutics, and biotechnological innovations.

## 5 Data availability

We have made all sequences, MSAs and predicted structures for Datasets 1–3 publicly available, along with the extracted RNA chains from references for Dataset 3. Additionally, we provide the list of PDB IDs for RNAs included in Dataset 4, and for complexes, we provide the RNA chains extracted from complex predictions and job files for the AlphaFold 3 web server that were used to obtain these predictions. All data can be accessed at https://zenodo.org/records/14561158.

The Protein Data Bank, from which we downloaded RNA structures and selected some for the creation of Dataset 3, is accessible at https://www.rcsb.org.

The AlphaFold 3 inference code was downloaded in November 2024 from https://github.com/google-deepmind/AlphaFold3 and was used for single-chain RNA predictions, while the web server, which can be accessed at https://alphafoldserver.com/welcome, was used for predictions of full complexes.

DRfold was downloaded in May 2023 as a standalone program from https://zhanggroup.org/DRfold/. It is also available as a web server and on GitHub at https://github.com/leeyang/DRfold.

DeepFoldRNA was downloaded in May 2023 from https://github.com/robpearc/DeepFoldRNA. It is also accessible at https://zhanggroup.org/DeepFoldRNA/ as both a web server and a standalone program.

RhoFold was obtained in May 2023 from a GitHub repository that is no longer available (https://github.com/RFOLD/RhoFold.git). However, an alternative repository is accessible at https://github.com/Dharmogata/RhoFold. RhoFold is also available as a web server at https://proj.cse.cuhk.edu.hk/aihlab/rhofold/.

RoseTTAFold2NA was obtained in June 2023 from https://github.com/uw-ipd/RoseTTAFold2NA. trRosettaRNA was downloaded in June 2023 from https://yanglab.qd.sdu.edu.cn/trRosettaRNA/download/. It is also accessible as a web server at https://yanglab.qd.sdu.edu.cn/trRosettaRNA/.

ARES was retrieved in July 2023 from https://zenodo.org/records/6893040.

The Rosetta suite, which provides Rosetta score, was downloaded in April 2024 from https://rosettacommons.org/software/download/.

USalign was acquired in May 2023 from https://zhanggroup.org/US-align/.

OpenStructure was utilized via a Docker container, which was downloaded in April 2024 from https://openstructure.org/download/.

For clash score calculations, we used the MolProbity package included in Phenix, downloaded in April 2024 from https://phenix-online.org/download/.

## 6 Supplementary data

Supplementary Data are available at NAR Online.

## 7 Competing interests

No competing interest is declared.

## 8 Author contributions statement

M.Š. and Y.Z. conceived the project and designed the experiments with the help of T.V. and Y.L. M.Š. and B.H. supervised the work. I.M. conducted the experiments, prepared data, and wrote the manuscript. T.V. participated in discussions and wrote the manuscript. All authors reviewed the manuscript and approved the final version.

## 9 Acknowledgments

The authors would like to thank Robin Pearce for the help with running DeepFoldRNA and valuable comments on this work, and Rafael Josip Penić on fruitful discussion on this work.

## Funding

This work was supported in part by the National Research Foundation (NRF) Competitive Research Programme (CRP) under Project *Identifying Functional RNA Tertiary Structures in Dengue Virus* [NRF-CRP27-2021RS-0001 to M.Š.]; in part by Agency for Science, Technology and Research (A*STAR) Industry Alignment Fund - Pre-positioning Programme (IAF-PP) under Project *Drugging RNA: Our Goal for Oncology (DRAGON)* [H23J2a0094 to M.Š.]; and in part by the Croatian Science Foundation under Project *Deep Learning-Based RNA Tertiary Structure Prediction and Generation* [IP-2024-05-1554 to M.Š.].

The computational work for this article was partially performed on resources of the National Supercomputing Centre, Singapore (https://www.nscc.sg).

## Competing interests

The authors declare no competing interests.

## Notes

### Competing Interest Statement

The authors have declared no competing interest.

### Summary of Updates

A major update of the manuscript

