## Supplementary for "A Comparative Review of Deep Learning Methods for RNA Tertiary Structure Prediction"

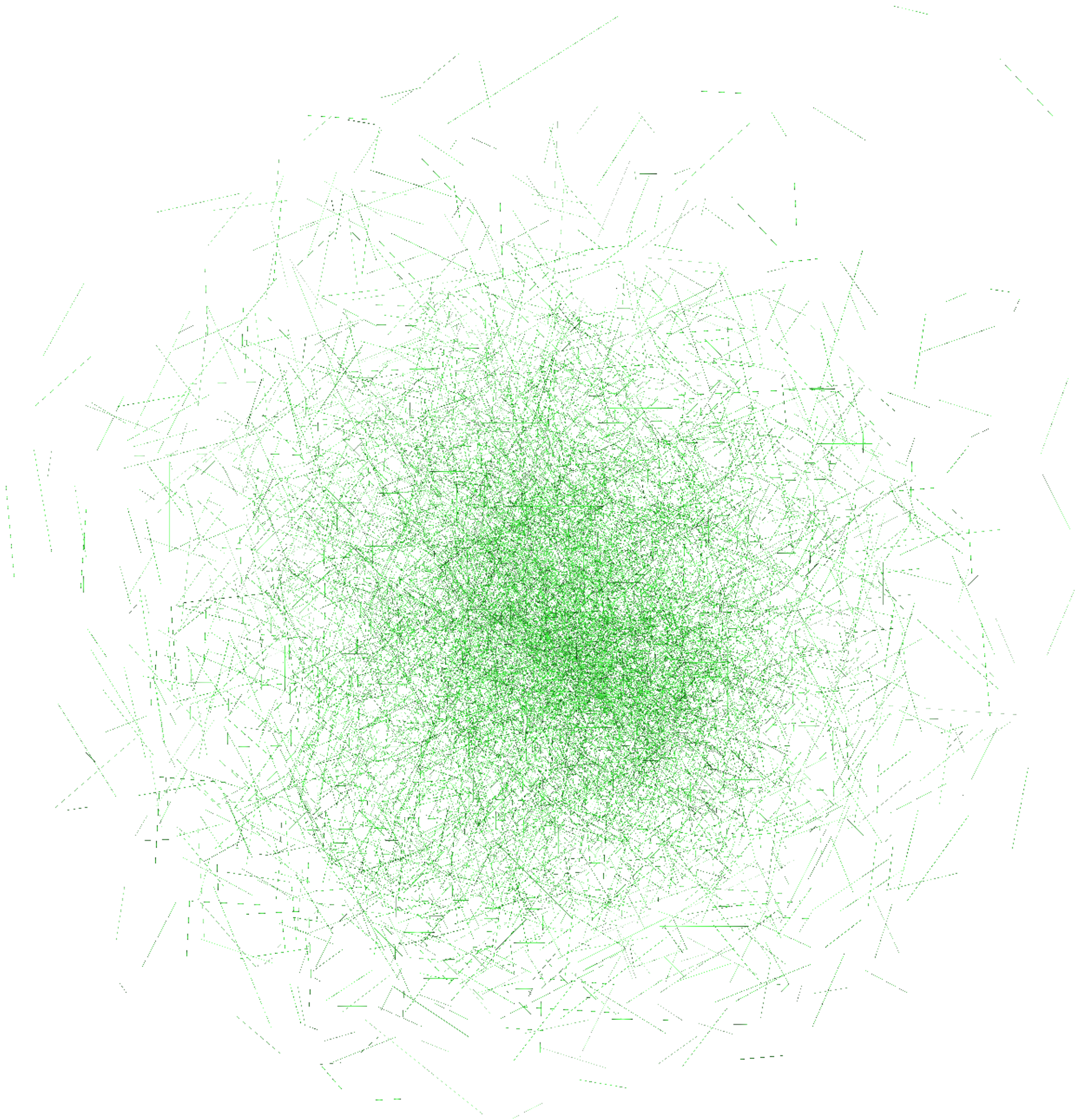

Figure S1: **RoseTTAFoldNA's prediction of complex with PDB_ID 8GU6. In this complex, there are, in total,** $2,488$ **residues.**

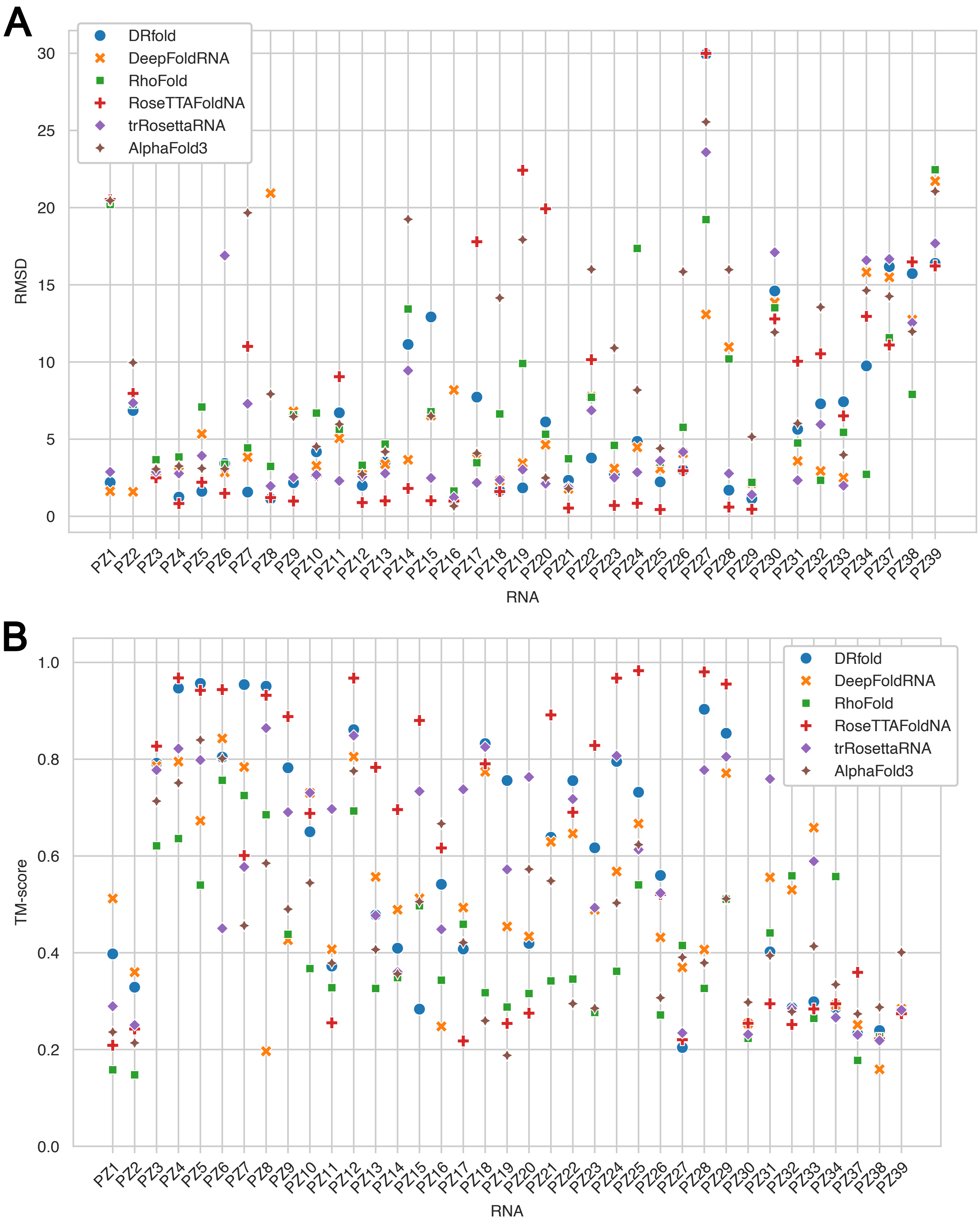


Figure S2: **All-atom RMSD and TM-score per RNA from Dataset 1 (RNA Puzzles)**. A. All-atom RMSD per RNA Puzzle. B. TM-score per RNA Puzzle.

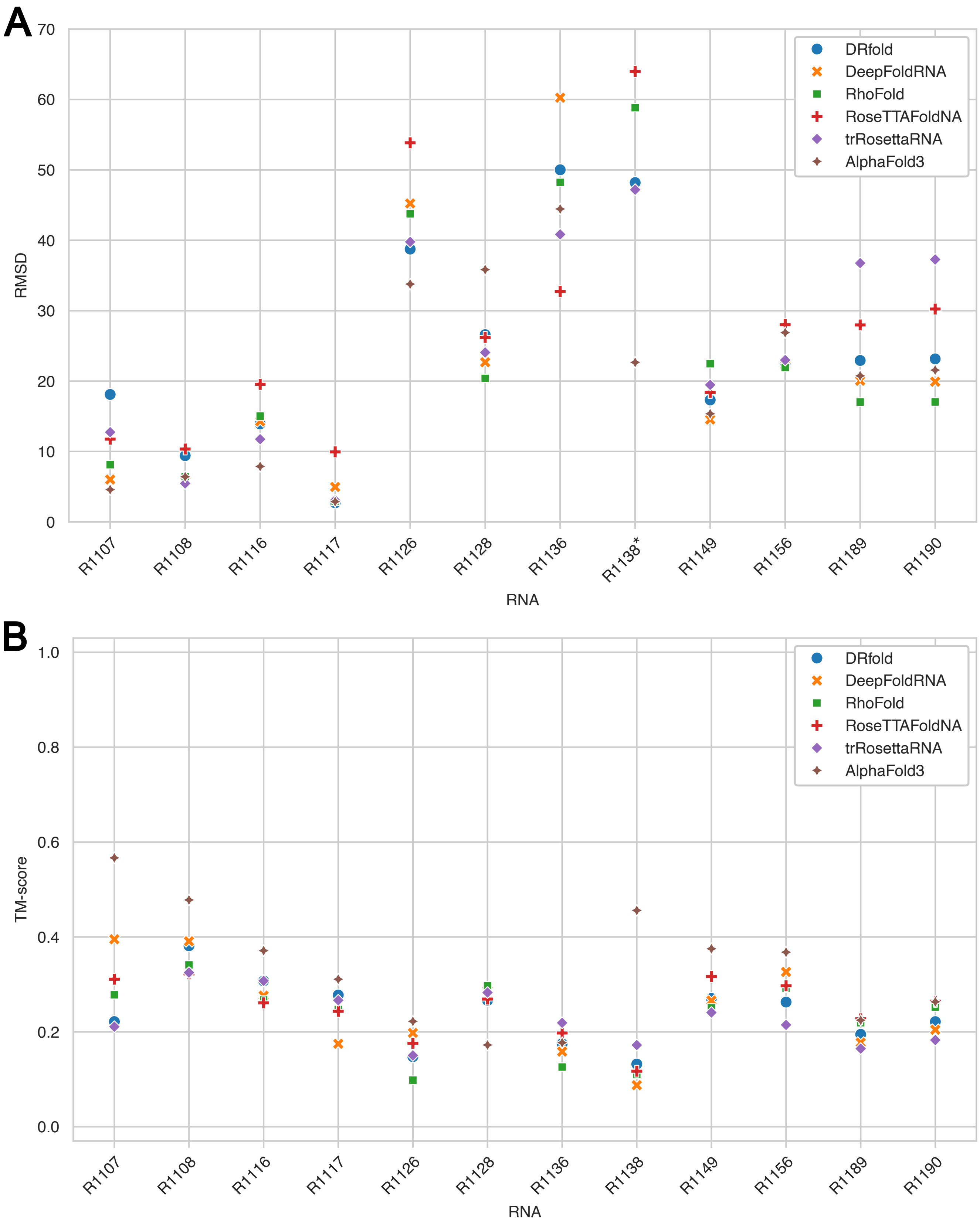


Figure S3: **All-atom RMSD and TM-score per RNA from Dataset 2 (CASP15 RNA target)**. A. All-atom RMSD per RNA target. B. TM-score per RNA target.

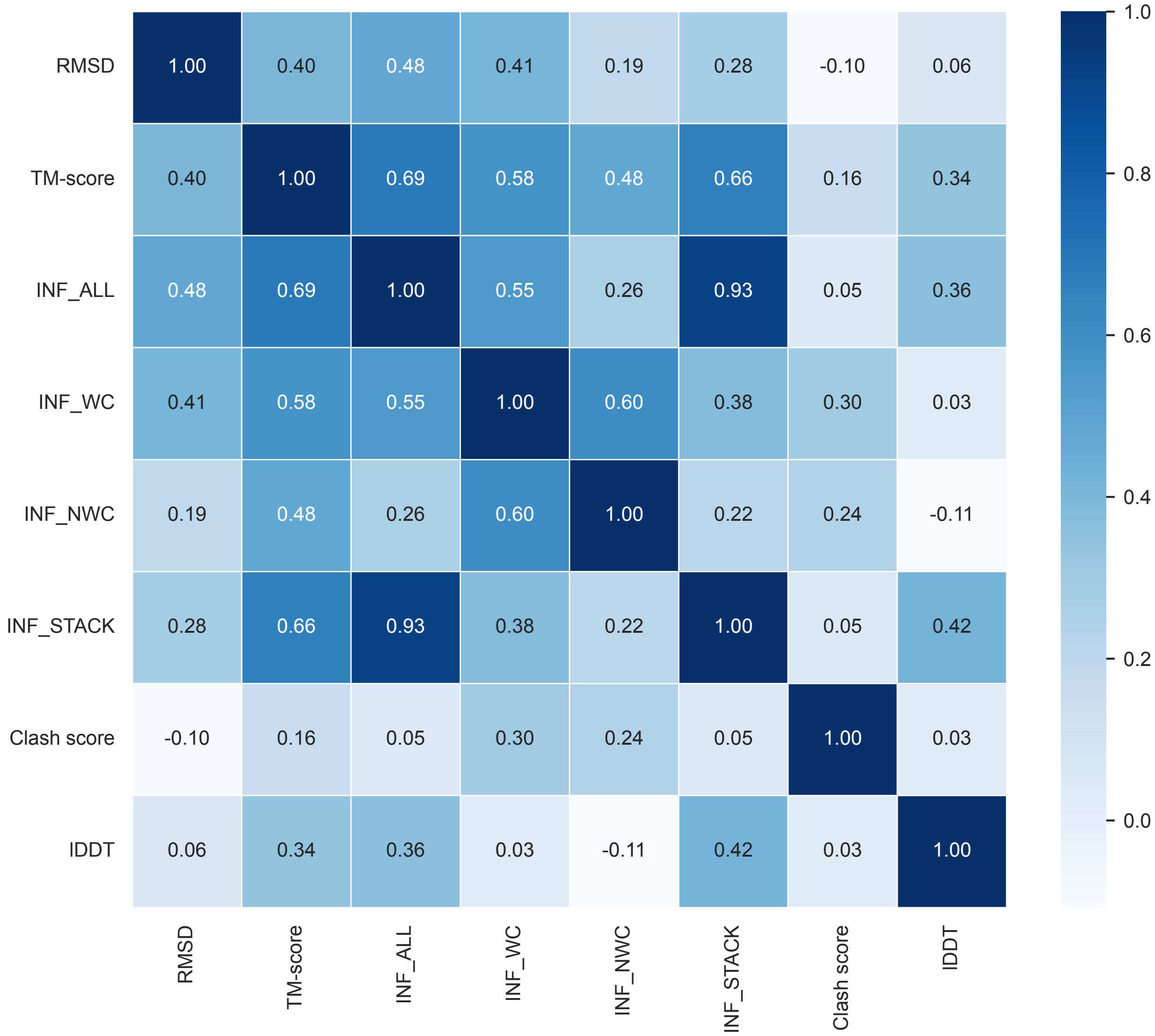


Figure S4: **Spearman correlation among different metrics for Dataset 3**. All-atom RMSD and clash score were reversed (negative value was taken) so that higher values uniformly represent better outcomes.

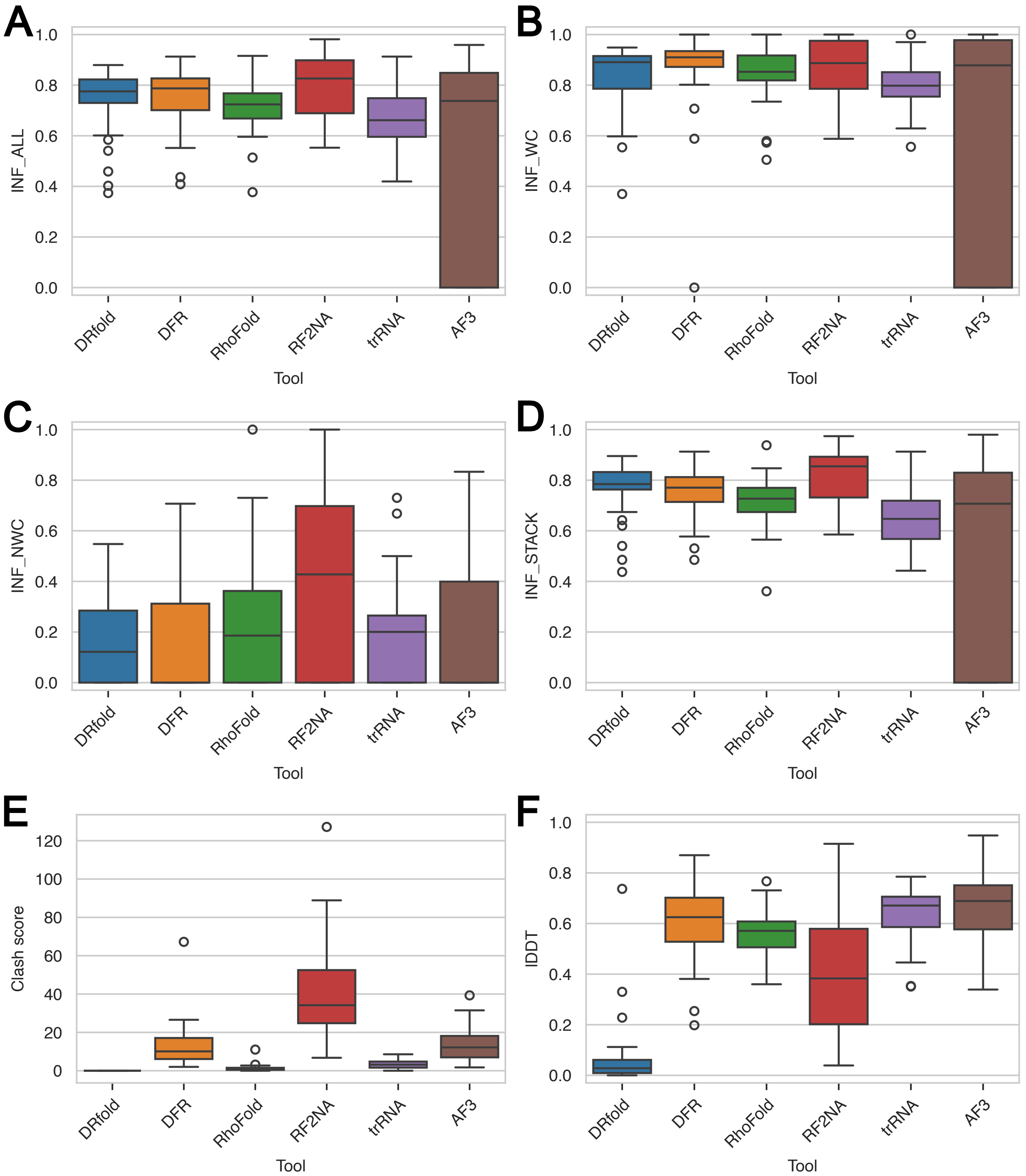


Figure S5: **Other important metrics for Dataset 1 (RNA Puzzles)**. A. INF_ALL (all RNA Puzzles have some interactions). B. INF_WC (excludes two RNA Puzzles without any Watson-Crick interactions). C. INF_NWC (excludes three RNA Puzzles without any non-Watson-Crick interactions). D. INF_STACK (all RNA Puzzles have stacking interactions). E. Clash score. F. lDDT. *Abbreviations used: DFR = DeepFoldRNA, RF2NA = RoseTTAFoldNA, trRNA = trRosettaRNA, AF3 = AlphaFold3.*

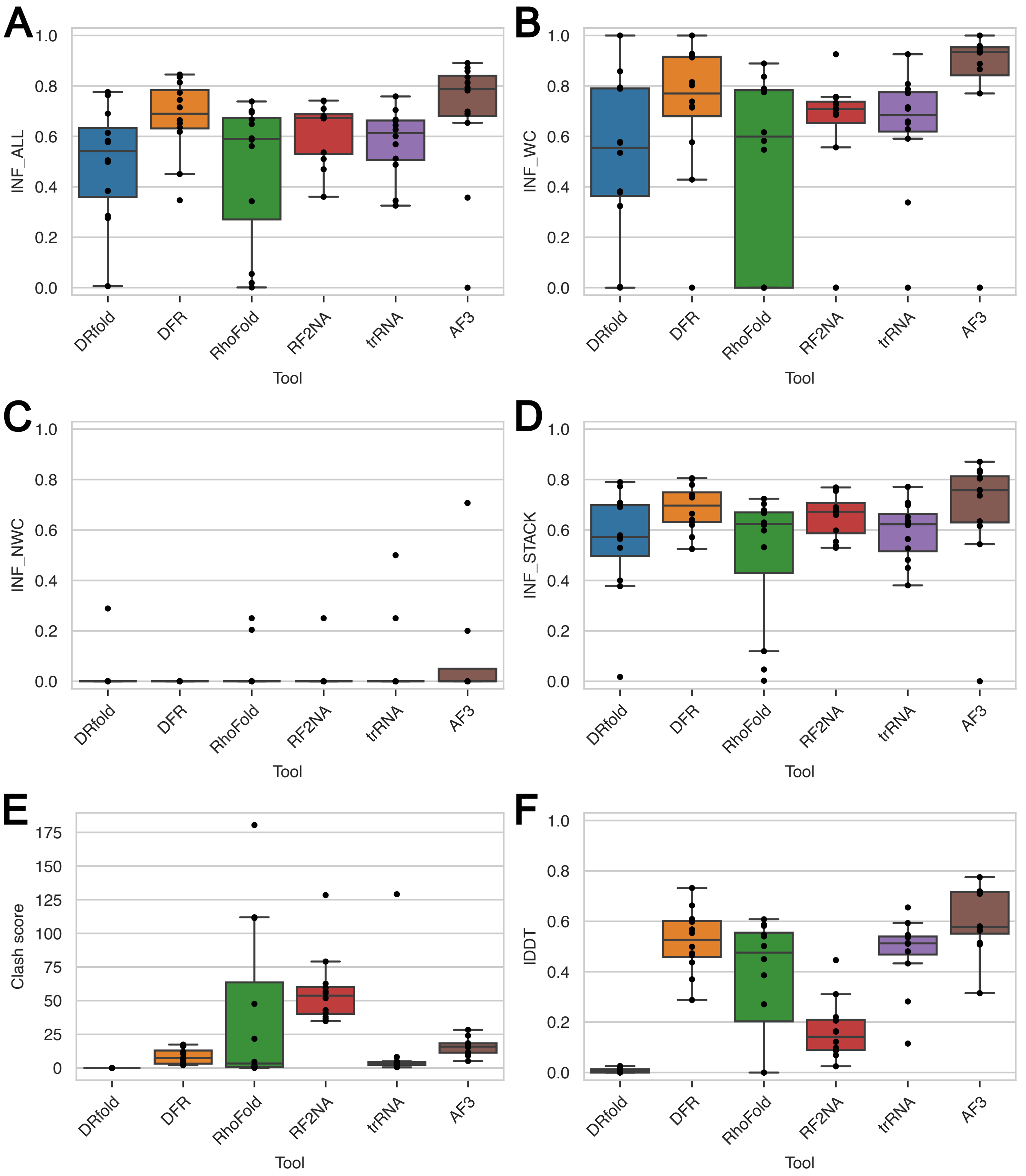


Figure S6: **Other important metrics for Dataset 2 (CASP15 RNA targets)**. A-D. All 12 RNA targets have all types of interactions (Watson-Crick, non-Watson-Crick and stacking interactions). A. INF_ALL. B. INF_WC. C. INF_NWC. D. INF_STACK. E. Clash score. F. lDDT. *Abbreviations used: DFR = DeepFoldRNA, RF2NA = RoseTTAFoldNA, trRNA = trRosettaRNA, AF3 = AlphaFold3.*

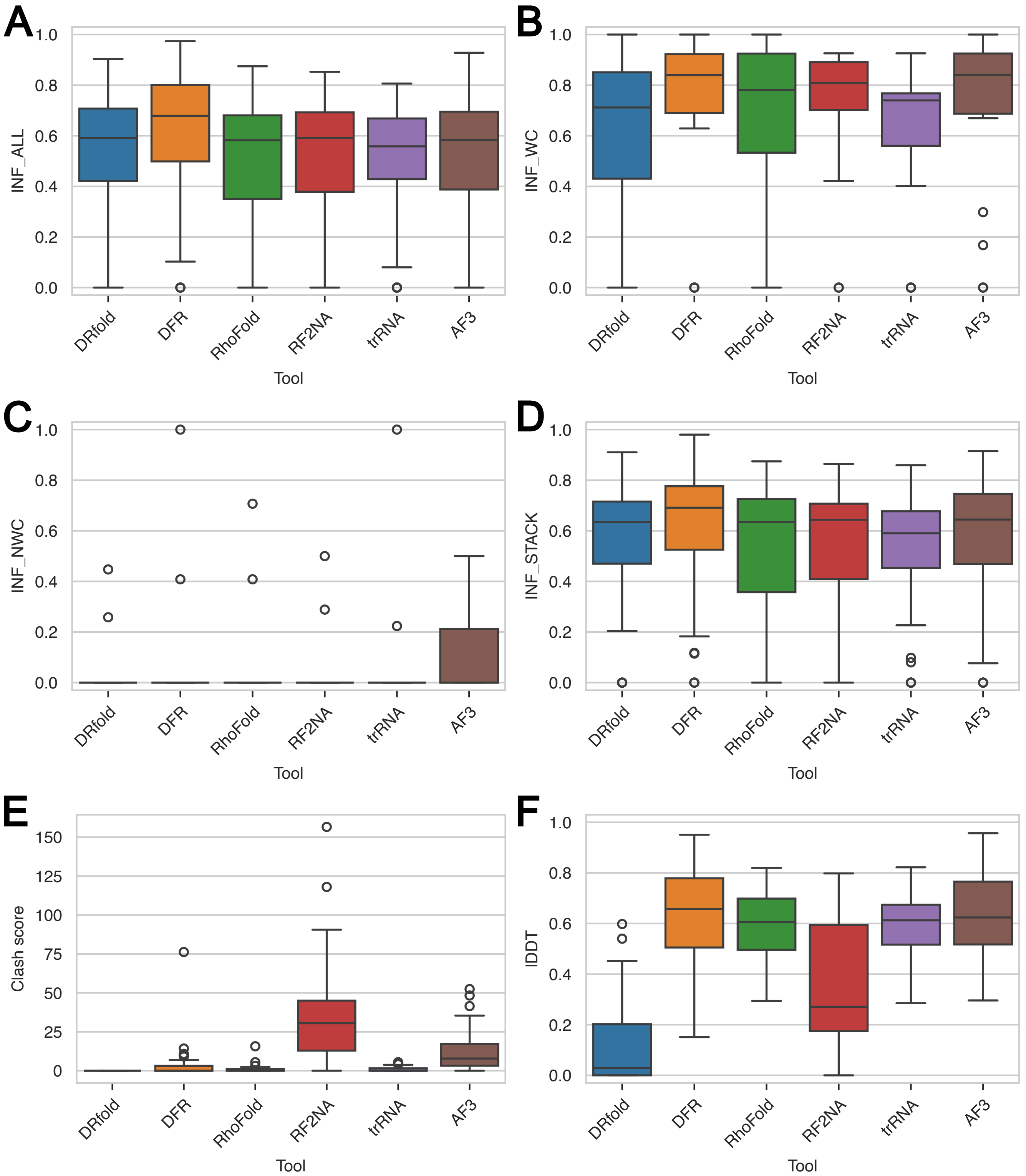


Figure S7: **Other important metrics for Dataset 4**. Dataset 4 is a subset of Dataset 3 where we did both structural and sequential clustering. A. INF_ALL (all RNAs have some interactions). B. INF_WC (excludes 42 RNAs without any Watson-Crick interactions). C. INF_NWC (excludes 45 RNAs without any non-Watson-Crick interactions). D. INF_STACK (all RNAs have stacking interactions). E. Clash score. F. lDDT. *Abbreviations used: DFR = DeepFoldRNA, RF2NA = RoseTTAFoldNA, trRNA = trRosettaRNA, AF3 = AlphaFold3.*

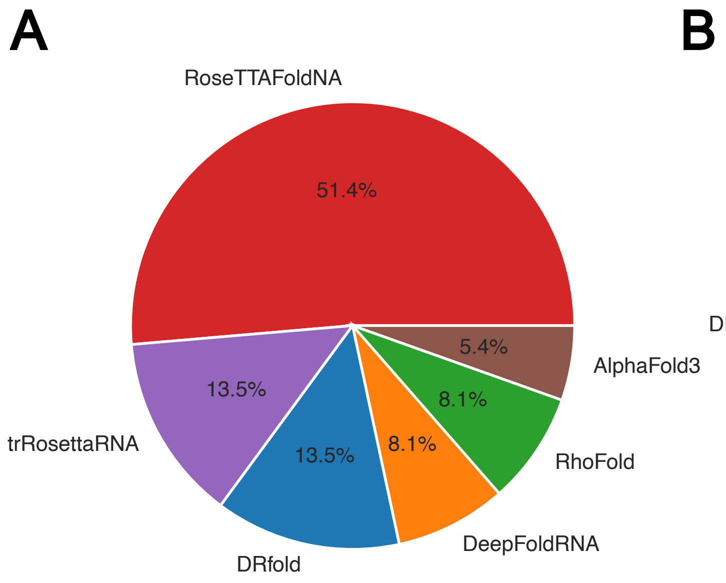


Figure S8: **Percentage of RNA chains for which each tool predicted the best structures according to lowest RMSD and highest TM-score for Dataset 1 (RNA Puzzles).** A. The best structure for each RNA is determined according to all-atom RMSD. B. The best structure for each RNA is determined according to TM-score.

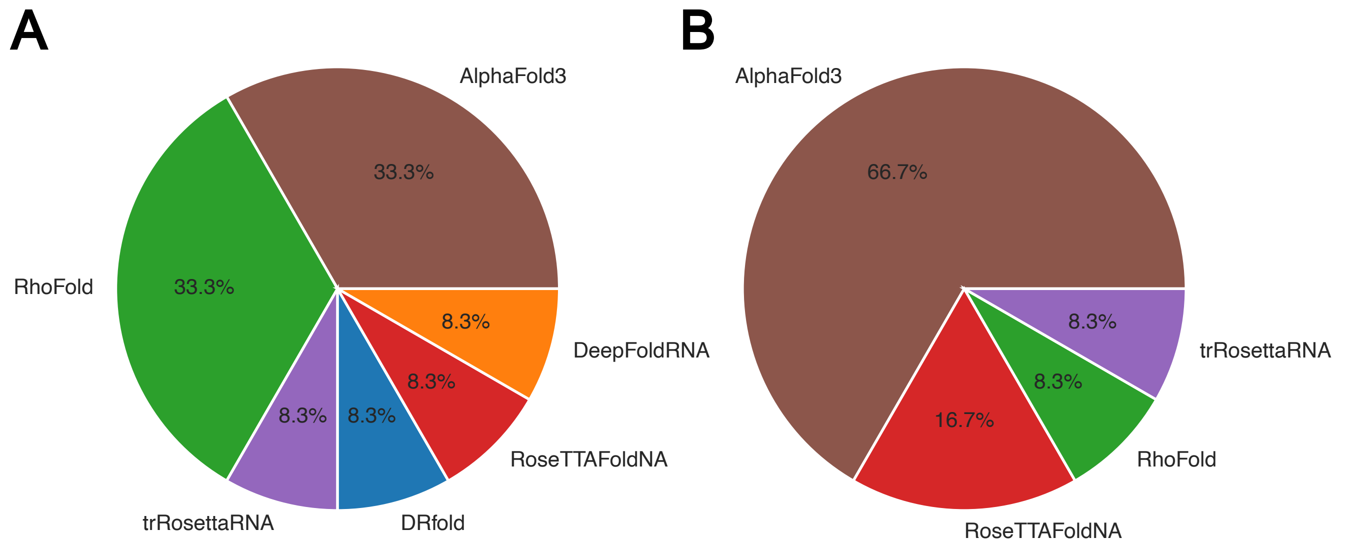


Figure S9: **Percentage of RNA chains for which each tool predicted the best structures according to lowest RMSD and highest TM-score for Dataset 2 (CASP15 RNA targets).** A. The best structure for each RNA is determined according to all-atom RMSD. B. The best structure for each RNA is determined according to TM-score.

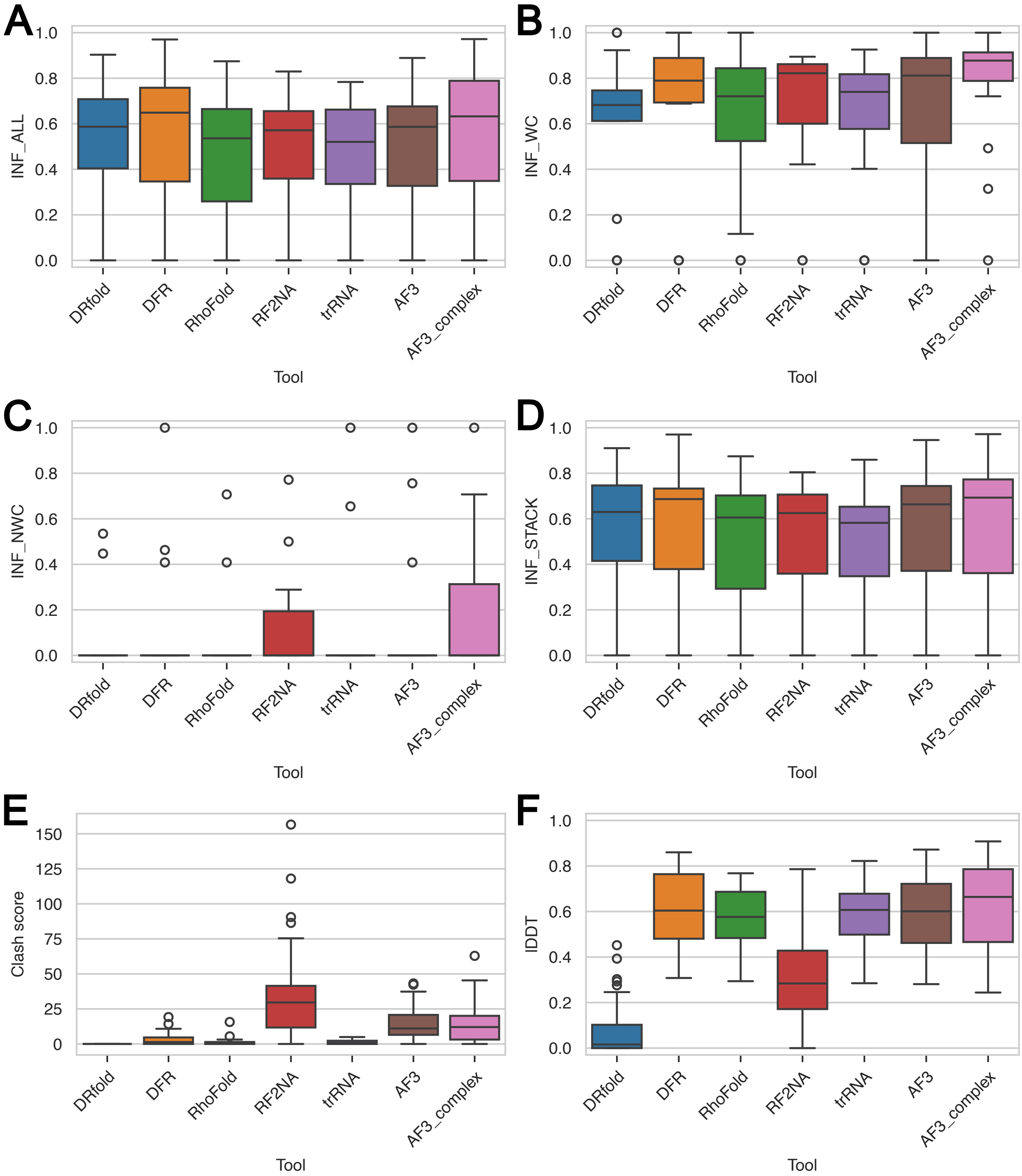


Figure S10: **Other important metrics for 43 RNA chains from Dataset 3 that can be predicted by AlphaFold3 as a complex (their complexes have less than 5000 residues).** The suffix '_complex' indicates predictions for the entire complex, not just the RNA chain within it. A. INF_ALL (all RNAs have some interactions). B. INF_WC (excludes 26 RNAs without any Watson-Crick interactions). C. INF_NWC (excludes 29 RNAs without any non-Watson-Crick interactions). D. INF_STACK (all RNAs have stacking interactions). E. Clash score. F. lDDT. *Abbreviations used: DFR = DeepFoldRNA, RF2NA = RoseTTAFoldNA, trRNA = trRosettaRNA, AF3 = AlphaFold3.*

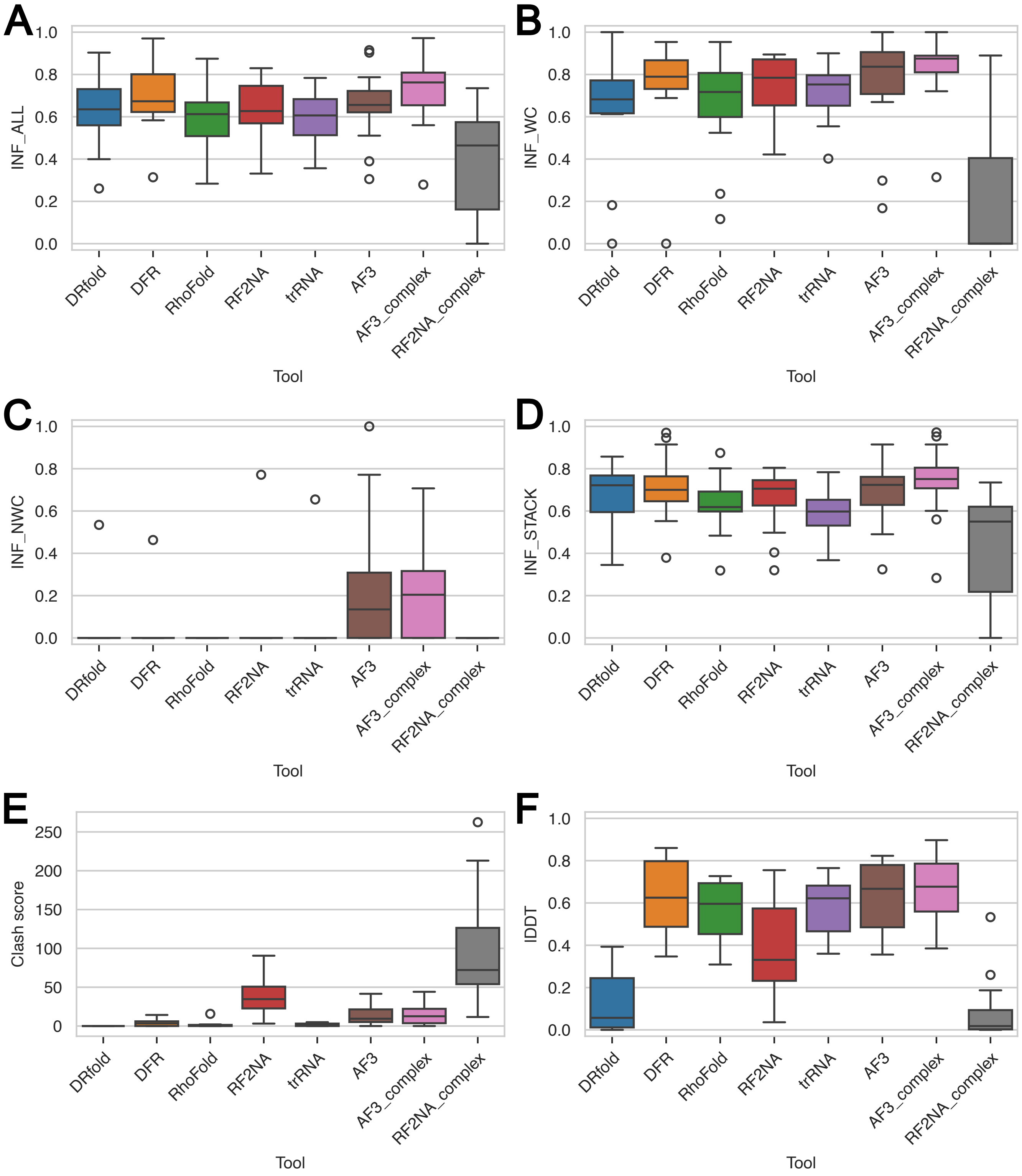


Figure S11: **Other important metrics for 19 RNA chains from Dataset 3 that were predicted by both RoseTTAFoldNA and AlphaFold3 as a complex**. The sufix '_complex' indicates predictions for the entire complex, not just the RNA chain within it. A. INF_ALL (all RNAs have some interactions). B. INF_WC (excludes 8 RNAs without any Watson-Crick interactions). C. INF_NWC (excludes 10 RNAs without any non-Watson-Crick interactions). D. INF_STACK (all RNAs have stacking interactions). E. Clash score. F. lDDT. *Abbreviations used: DFR = DeepFoldRNA, RF2NA = RoseTTAFoldNA, trRNA = trRosettaRNA, AF3 = AlphaFold3.*

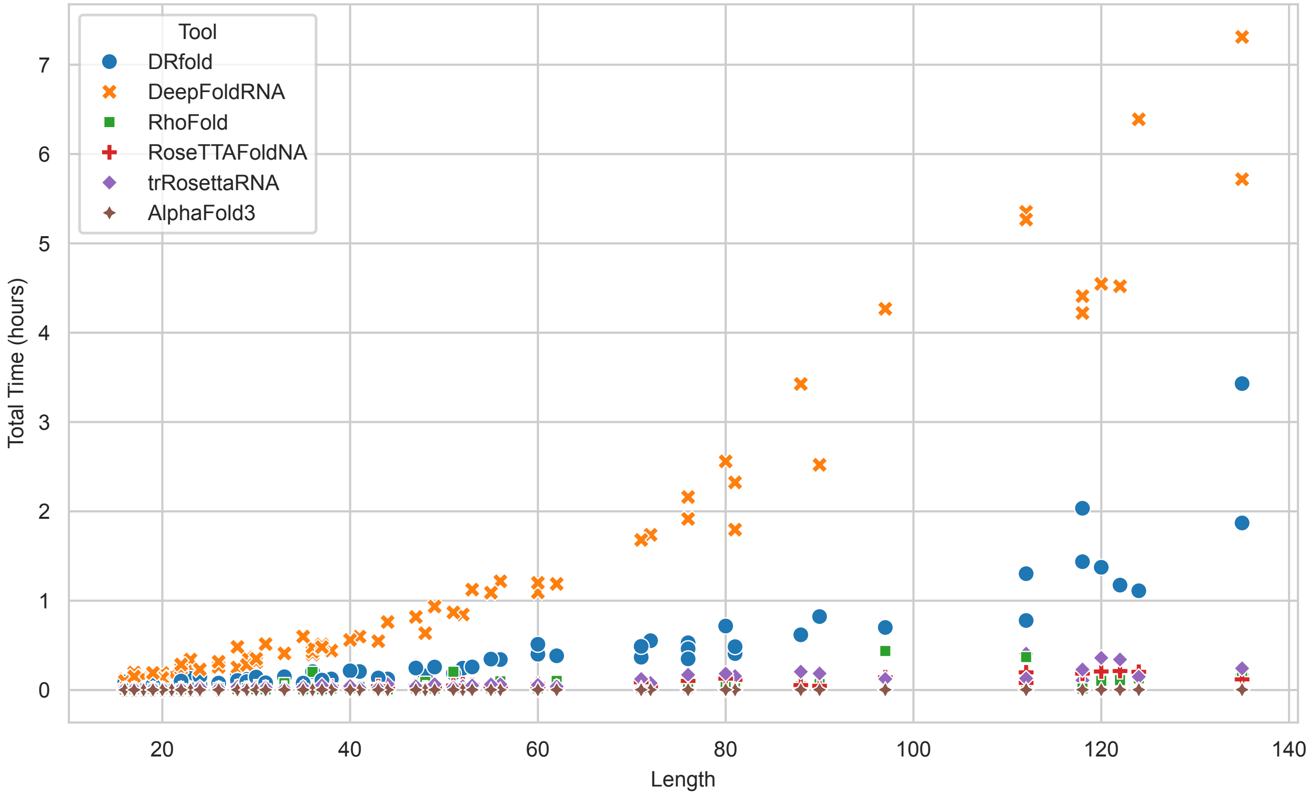


Figure S12: **Execution time for RNAs from Dataset 3.**
